# Structures of *Chaetomium thermophilum* TOM complexes with bound preproteins

**DOI:** 10.1101/2025.04.16.649081

**Authors:** Ahmed-Noor A. Agip, Pamela Ornelas, Tzu-Jing Yang, Ermanno Uboldi, Sabine Häder, Melanie A. McDowell, Werner Kühlbrandt

## Abstract

Mitochondria import most of their proteins from the cytoplasm through the TOM complex. Preproteins containing targeting signals are recognized by the TOM receptor subunits, and translocated by Tom40 across the outer mitochondrial membrane. We present four structures of the preprotein-bound and preprotein-free TOM core and holo complex from the thermophilic fungus *Chaetomium thermophilum*, obtained by single-particle electron cryomicroscopy. Our structures reveal the symmetric arrangement of two copies of the Tom20 receptor subunit in the TOM holo complex. Several different conformations of Tom20 within the TOM holo complex highlight the dynamic nature of the receptor. The structure of preprotein-bound Tom20 provides insight into the early stages of protein translocation.

## Introduction

Mitochondria are central to energy metabolism and other important physiological processes that require the import of approximately 1,000 different proteins (1). The imported proteins carry a mitochondrial targeting signal (MTS) that leads them to their respective locations within the organelle (2). The translocase of the outer mitochondrial membrane (TOM) is the entry gate for the majority of proteins imported into mitochondria as precursor proteins, or preproteins. In humans and fungi, the TOM core complex is a membrane-embedded dimer of twofold symmetry, consisting of two copies each of the β-barrel Tom40 pore, the α-helical Tom22 receptor, and the small α-helical structural subunits Tom5, Tom6, and Tom7 (3). Two other preprotein receptors, Tom20 and Tom70, are loosely bound to the TOM core, forming the TOM holo complex (4). Tom22 and Tom20 both recognize the MTS within N-terminal presequences, whereas Tom70 recognizes internal MTS such as those present in the mitochondrial carrier proteins (5, 6).

The first cryoEM structures of the TOM complex were determined from *Saccharomyces cerevisiae* at 18 Å resolution (7), and later from *Neurospora crassa* at 6.8 Å (8). The *N. crassa* structure revealed the general architecture of the TOM complex and its subunits, including two Tom40 pores tilted at an angle of 20° relative to the membrane normal, bridged by two Tom22s and surrounded by the small Tom5-7 subunits (8). Subsequent cryoEM structures of the TOM core complex all conformed to the same architecture (9–16). The structure of the TOM holo complex has been studied more recently. The structure of a cross-linked human TOM complex shows two copies of the Tom20 receptor domain on the cytoplasmic face of the complex (16). Structures of the native *N. crassa* (14) and *Homo sapiens* TOM show one resolved copy of Tom20 per dimer in two different conformations (17). The Tom70 receptor has additional roles beyond protein translocation, including the formation of ER contact sites and the recognition of viral factors (18, 19) but seems to be loosely attached to the complex. The structure of the Tom70 soluble domain has been determined (18), but its complete structure in association with the TOM complex remains unresolved.

Although the overall pathway for mitochondrial protein translocation is known, specific interactions of the TOM subunits with preproteins, as well as conformational changes that enable translocation, are not well-characterized. It has been shown that downstream interaction partners, including the translocase of the inner membrane (TIM) complexes TIM22 and TIM23, are required for complete translocation (20–23). However, we do not understand how preproteins engage with TOM.

In this work, we take a step towards understanding the translocation mechanism of TOM through purification and structure determination of intact, thermostable complexes from the thermophilic fungus *Chaetomium thermophilum*. We determined TOM core and holo structures with and without the bound MTS of rat aldehyde dehydrogenase (pALDH), revealing numerous interaction sites with the Tom20 receptor and the Tom40 translocation pore.

The improved stability of the *C. thermophilum* complex yielded a 2.7 Å resolution map of the core complex, enabling us to observe high-resolution features, in particular bound lipids and detergent molecules, indicating that lipid binding sites are conserved across species. Likewise, in our 3.2 Å resolution structure of the symmetric TOM holo complex, two copies of the Tom20 receptor interact with each other at their receptor domains and are stabilized by the cytoplasmic domain of Tom22, consistent with our previous *N. crassa* model. Additionally, we identified an idle resting state and a continuous movement of Tom20 in the early stages of preprotein translocation.

## Results

### Purification and cryoEM preparation of TOM complex from *Chaetomium thermophilum*

We decided to isolate the *C. thermophilum* TOM complex for structure determination, as protein complexes from this thermophilic fungus have previously been shown to have a superior stability and rigidity for cryoEM (24). To do this, we integrated a copy of Tom22 with a C-terminal FLAG epitope into the genome of *C. thermophilum*. We first extracted the mitochondrial complexes after solubilization with the detergent glyco-diosgenin (GDN). After affinity purification and size-exclusion chromatography, we obtained a single monodisperse peak containing *C. thermophilum* TOM **(Fig. S1A)**. CryoEM grids prepared with the purified sample had an optimal particle distribution **(Fig. S1B)**. In order to understand the molecular interactions of TOM with preproteins, we further purified TOM and incubated it with the presequence of the well-characterized rat pALDH preprotein before cryoEM analysis (14).

We carried out cryoEM image processing separately for TOM prepared with or without pALDH. By performing iterative rounds of 2D and 3D classifications, we obtained a C2-symmetrized map of the TOM core complex at 2.7 Å resolution for the pALDH-bound state **(Fig. S2)**. Applying 3D variability analysis to the same set of particles resulted in two unique classes of the TOM holo complex that exhibited a major connecting density above the cytoplasmic face of the Tom40 twin pores. Joint refinement of these two classes resulted in a C2-symmetrical map of the TOM holo complex at 3.2 Å resolution **(Fig. S2)**. In the preparation without pALDH, iterative rounds of particle sorting produced reconstructions of both the TOM core and holo complex at 3.2 Å and 3.8 Å resolution, respectively **(Fig. S3)**. Visual comparison of the corresponding maps from both preparations revealed no differences **(Fig. S4)**, indicating that incubation of the isolated complex with pALDH does not induce major conformational changes. For this reason, and due to their significantly higher resolution, we focus primarily on the pALDH-bound TOM complexes, noting that any interpretation of the structures may equally apply to the complex without bound preprotein.

### Structure of the TOM core complex

At 2.7 Å global resolution, we were able to build the core components of the TOM complex with confidence **(Table S1, Fig. S5)**. As expected, the TOM core complex conforms to the same composition and architecture as previous structures from other species **(Fig. S6)** (8–16). In brief, the complex is composed of the central Tom40 twin pore, the main receptor Tom20 and the small structural subunits Tom5-7 (**Figure 1A-B**). With the exception of Tom40, which is a nineteen-strand β-barrel, all other subunits have a single transmembrane helix. As with other species, the dimeric complex is shaped like a shallow funnel, where the protomers are tilted towards each other at the cytoplasmic side, and contact between the two Tom40s is mediated through β-strands 1, 2 and 19. On the intermembrane space (IMS) side, the barrels are separated by two Tom22 helices and a central lipid (**Figure 1C**). At the protomer interface, the Tom22-lipid interaction causes the complex to tilt by 20° relative to the membrane normal, as previously reported for *N. crassa* TOM (8, 14). Whether the funnel shape of the complex is of functional significance for the translocation process is currently unknown.

**Figure 1.**
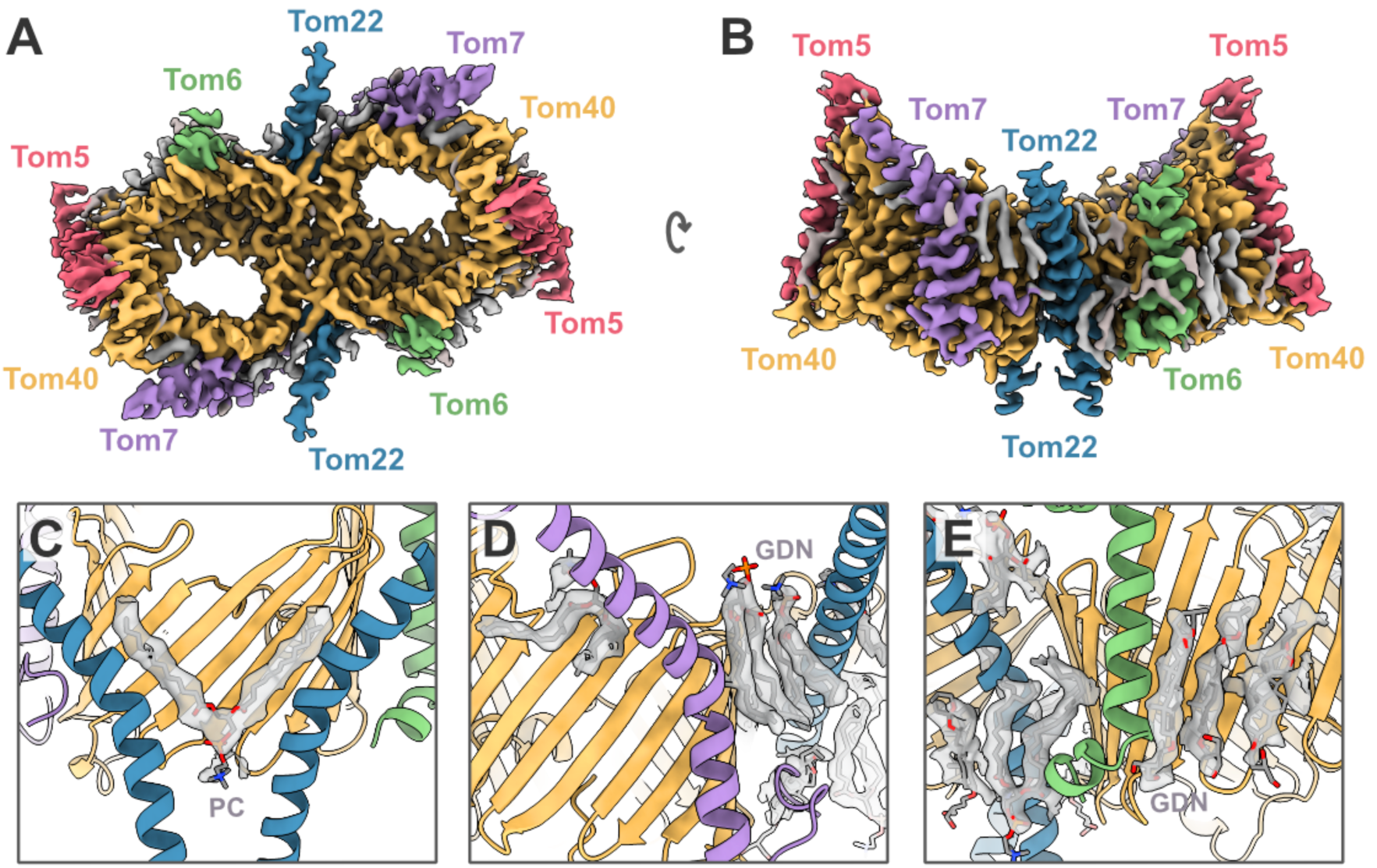
*C. thermophilum* TOM core complex. **(A)** CryoEM map of the TOM core complex dimer at 2.7 Å resolution, colored by subunit: Tom40, yellow; Tom22, blue; Tom5, Tom6 and Tom7 are pink, green and purple, respectively. Lipid and detergent molecules are grey. **(B)** Side view of the map along the plane of the outer mitochondrial membrane. **(C-E)** Protein-lipid and protein-detergent interactions surrounding the TOM core dimer. Lipids were modeled as PC and detergent molecules as GDN.

*C. thermophilum* Tom40 is formed of two N-terminal helices followed by nineteen consecutive β-strands and a C-terminal extension (**Fig. S7**). As with *N. crassa* and *S. cerevisiae, C. thermophilum* Tom40 helix α1 (absent in human TOM) starts at the IMS (10, 12, 14), then threads through Tom40, forming the pore-stabilizing helix α2, which finally emerges on the cytoplasmic side. Similar to *N. crassa*, the C-terminal helix α3, which we modeled as a random coil, re-enters the Tom40 pore. The remaining TOM subunits are very similar to the previously described structures (8–16), with the Tom22 helical region protruding into the IMS, Tom7 displaying its characteristic Z shape, the Tom6 N-terminal region being unresolved and Tom5 forming a well-defined helix at right angles to the membrane.

We identified a number of unassigned densities in our TOM core complex structure. Five in each protomer were modeled as phosphatidylcholine (PC) lipids, resulting in a total of eleven lipids in the dimeric complex, including the interfacial lipid. Additionally, we modeled four detergent molecules per protomer as GDN. Four of the lipids, one at the Tom40 interface and another three in each protomer, are observed in the same positions as in *N. crassa* and *H. sapiens*, indicating that their binding sites are conserved **(Fig. S8)**. The interfacial lipid is critical in bridging the two protomers at the IMM-cytoplasm interface (**Figure 1C**). Three of the conserved lipids are at the membrane-cytoplasm interface, with one and two lipids bound at the interface of Tom7-Tom40 and Tom7-Tom20-Tom40, respectively (**Figure 1D**). The two remaining non-conserved lipids are bordered by Tom20, Tom6 and Tom40, with their head groups facing in opposite directions (**Figure 1E**). Presumably, owing to the rigidity of Tom40, we observed several clusters of small unassigned densities within the barrel that could be interpreted as bound water molecules or ions **(Fig. S9)**. We note that these densities are present at the interface between helix α2 of Tom40 and the barrel, indicating the inner helix is stable and unlikely to be involved in the conformational changes important for gating preprotein translocation.

### Stoichiometry of Tom20 in the *C. thermophilum* TOM holo complex

Owing to the known flexibility of Tom20 (14), we focused on the cytoplasmic densities above the Tom40 pores. The analysis resulted in a 3.2 Å map of the TOM holo complex with two copies of Tom20 resolved **(Fig. S10)**. Consistent with our previous *N. crassa* structures, the core subunits exhibited the clearest features with a strong drop in resolution at the periphery and the Tom20 subunits (9–16).

In the structure, the Tom20 dimers appear to be symmetrical, even before imposing C2 symmetry during the last refinement step **(Fig. S2)**. The symmetric Tom20 receptors extend along the membrane plane and rest above the twin Tom40 pores (**Figure 2**). Tom20 does not adapt to the funnel curvature of the pore but maintains a position above and roughly parallel to the membrane. Tom20 consists of seven helices, α1-7 **(Fig. S7)**, of which α2-6 were clearly resolved. Consistent with the *N. crassa* TOM map, Tom20 emerges from the detergent micelle as it interacts with the N-terminus of Tom22 **(Fig. S11A)**. We propose an electrostatic interaction between Tom20 and Tom22 that enables the incorporation of Tom20 into the holo complex. Given this stabilizing interaction, we were able to model more residues towards the N-terminus of Tom22 compared to the core complex. Further, sequence alignment identified conserved charged residues that mediate contact **(Fig. S11B)**. We therefore posit that the binding of Tom20 to the complex is likewise conserved, at least in metazoa and fungi. To test our prediction, we used AlphaFold3 to generate a model of the human TOM holo complex for visual comparison **(Fig. S12)**(25). In this model, the interaction of Tom20 and Tom22 resembles that in our experimental structure of the *C. thermophilum* holo complex.

**Figure 2.**
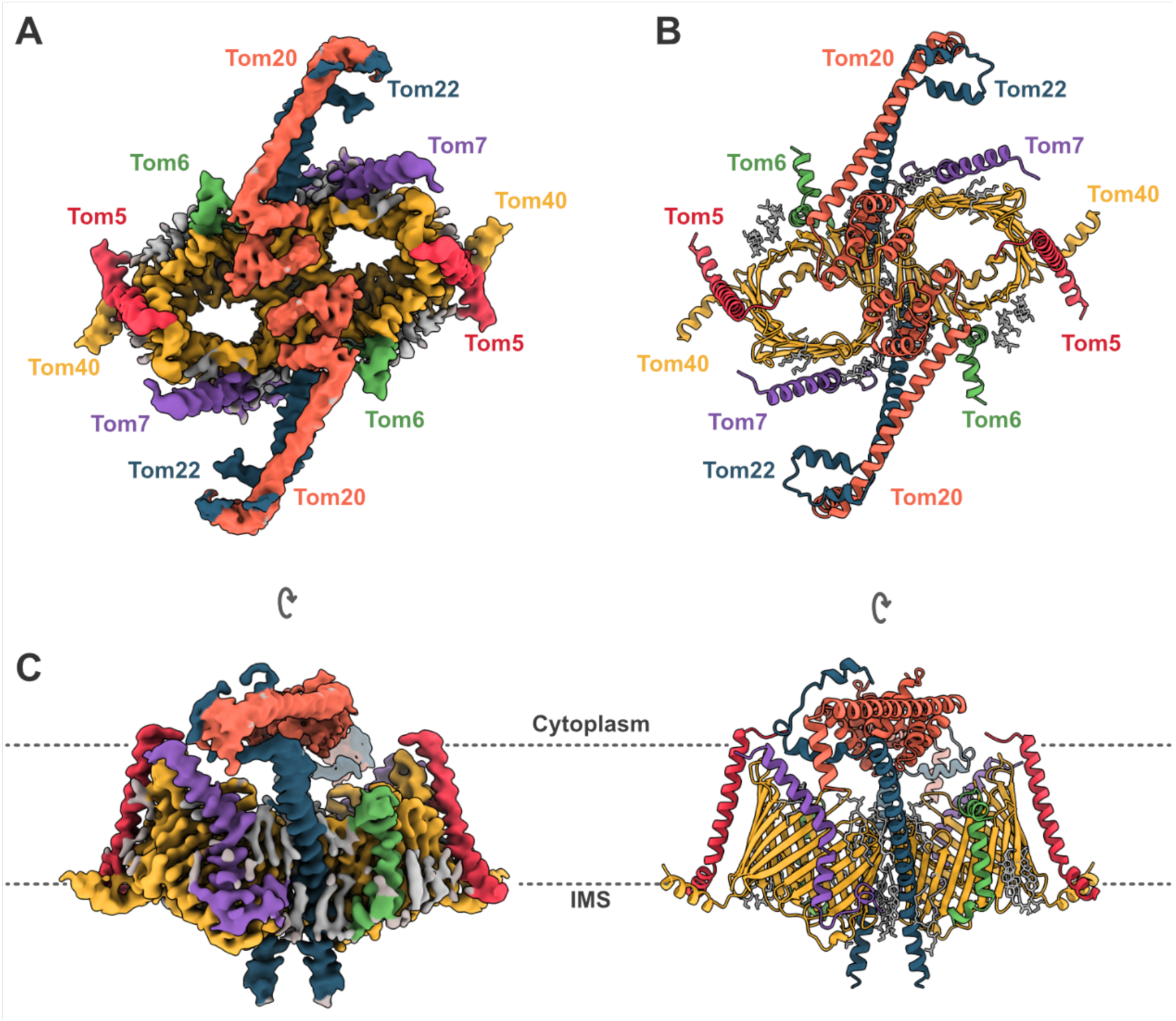
*C. thermophilum* TOM holo complex. CryoEM map and model of the TOM holo complex colored by subunit: Tom40, yellow; Tom22, blue; Tom20, orange; Tom5, Tom6 and Tom7 are red, green and purple, respectively. Lipid and detergent molecules are grey. (**A**) TOM holo complex dimer at 3.2 Å resolution viewed from the cytoplasm. (**B**) Cartoon representation of the corresponding atomic model. (**C**) Side view along the plane of the outer mitochondrial membrane. The lipid bilayer is indicated by dashed lines.

### Preprotein binding to *C. thermophilum* TOM

The receptor domain of Tom20 at the C-terminus of the protein includes a tetratricopeptide repeat (TPR) motif forming a hydrophobic pocket that binds to mitochondrial targeting sequences (26, 27). The superposition of our model with crystal structures of the rat Tom20 receptor domain (PDB 3AWR) indicated that they are almost identical, with an RMSD of less than 1 Å **(Fig. S13)**. On closer inspection, we found that the hydrophobic binding grooves of the Tom20 receptors are orientated towards each other. However, the contact surface area between the two receptors is small, measuring only 185 Å^2^. In addition, the local resolution at the Tom20-Tom20 contact is relatively low **(Fig. S10B)**, suggesting that this region of the protein is dynamic, as expected for a receptor that has to be mobile to initiate protein translocation.

Symmetry-relaxed refinement results in additional densities corresponding to the preprotein pALDH caught in mid-translocation inside both Tom40 pores near the hydrophobic pocket of the Tom20 TPR. We generated a difference map against the unbound TOM holo map that highlights the presence of the preprotein and its binding partners (**Figure 3 and Fig. S14)**.

**Figure 3.**
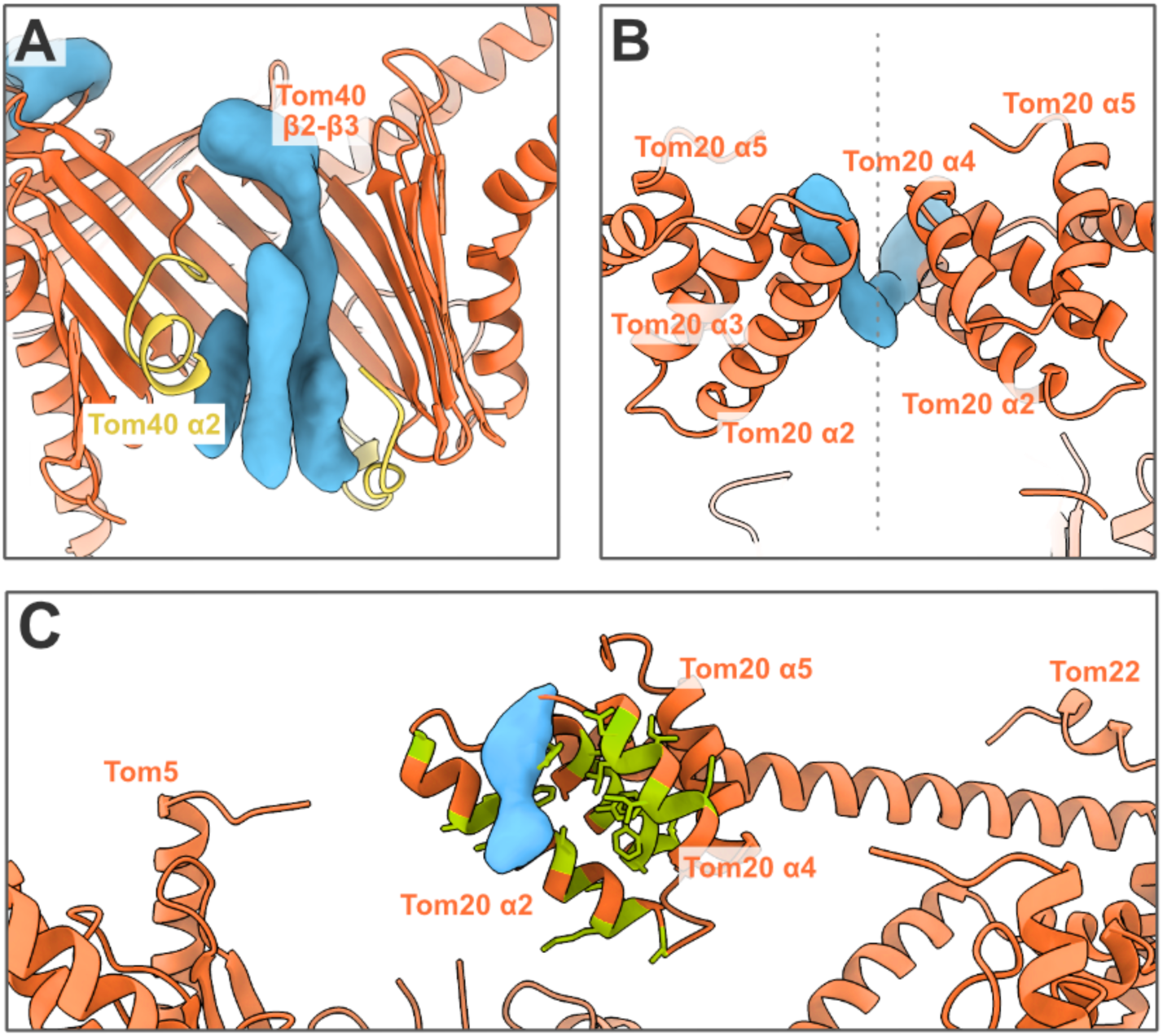
TOM holo model with preprotein densities in the cryoEM difference map. (**A**) Preprotein densities (blue) within the Tom40 translocation pore, delimited by the hydrophobic residue-rich inner Tom40 helices (yellow). (**B**) Tom20 receptor domains interacting with preprotein density (blue). The dashed line indicates the symmetry axis. (**C**) Close-up of one Tom20 receptor interacting with the preprotein density. Hydrophobic regions are shown in green.

In the difference map, three elongated densities occupy the space inside one of the Tom40 translocation pores, suggesting multiple preprotein binding sites (**Figure 3A**). The longest preprotein density interacts with the loop between Tom40 strands *β*2-*β*3 on the cytoplasmic side of the pore (**Fig. S7A)**, suggesting a possible entrance contact site, in accordance with previously published cross-linking studies (28). On the IMS side, the density makes contact with the hydrophobic C-terminal extension of Tom40. The remaining densities in the pore are shorter and delimited by the hydrophobic inner Tom40 helix, α2. On the cytoplasmic side, the preprotein appears in two positions as it interacts with each Tom20 receptor domain, between helices α4-α5 in one case, and α3 in the other (**Figure 3B and Fig. S7B**). **Figure 3C** shows a detailed view of one of these interactions, indicating the hydrophobic residues that line the receptor pocket. Both densities are close to the position of the preprotein in previously published crystal structures of the Tom20 receptor domain **(Fig. S13)** (26).

### Continuous flexibility of the Tom20 receptors

As seen in **Figure 2**, the twin Tom20 receptors meet at the center of the complex in what we propose to be a resting conformation. This conformation differs from those previously reported in *N. crassa* and *H. sapiens* **(Fig. S15)** (*14, 16*). Further evaluation by 3D classification of TOM without bound preprotein **(Fig. S3)** revealed that Tom20 adopts various conformations, consistent with our results on the *N. crassa* TOM (14). We discern four conformations of Tom20 in the absence of a preprotein. Following heterogeneous and non-uniform refinement of the resulting classes, we attained structures ranging from 4.4 to 5.1 Å resolution, showing Tom20 in different positions along what we assume to be a continuous trajectory (**Figure 4**).

**Figure 4.**
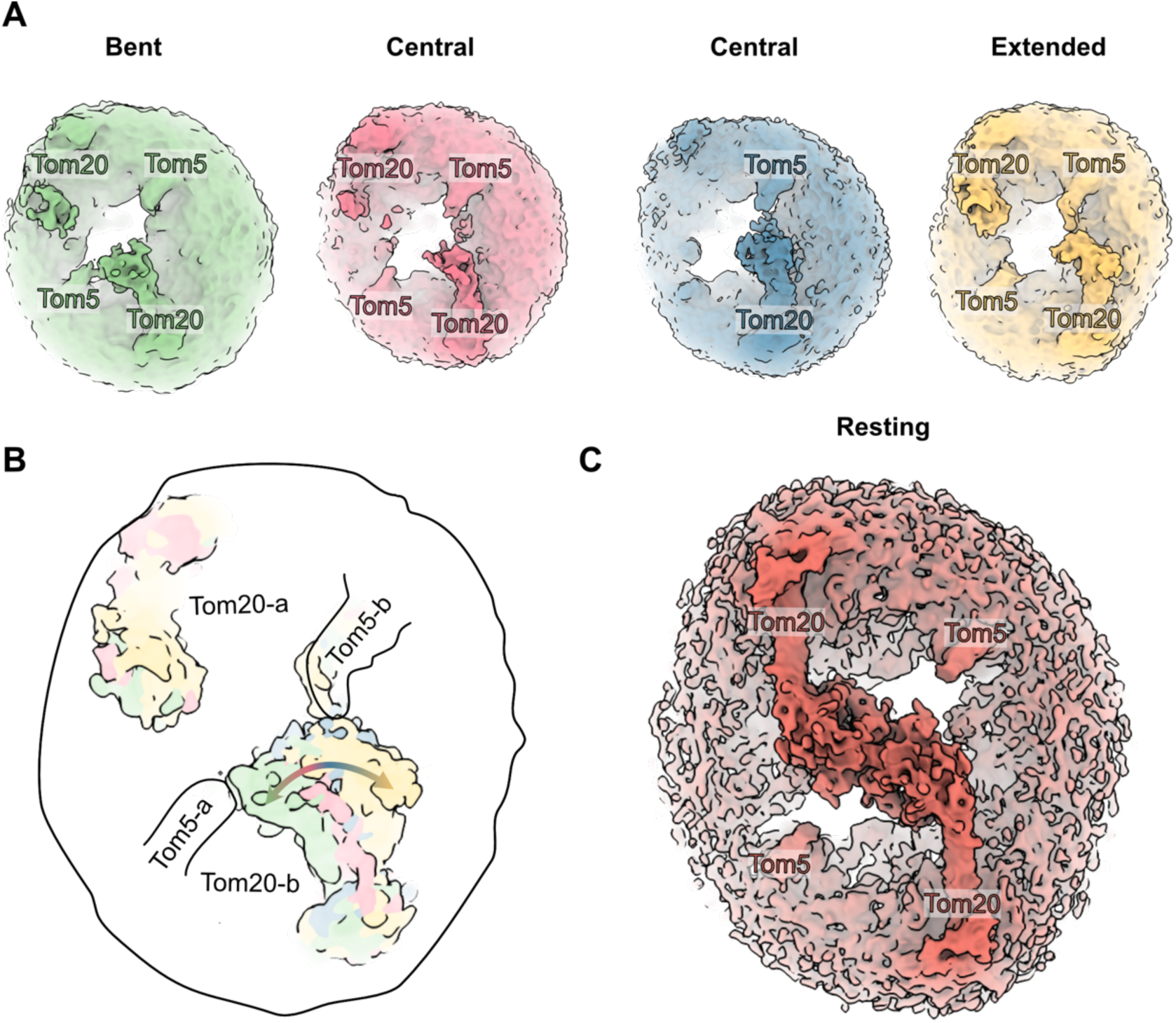
Tom20 takes on various conformations on the cytoplasmic membrane surface. (**A**) 3D classification indicates a range of Tom20 conformations from bent to central to extended, left to right. (**B**) A schematic representation of superposed TOM complexes with Tom20 adopting different conformations as it interacts with Tom5. a and b denote the individual protomers. (**C**) The resting conformation shows the twin Tom20 receptors interacting with one another, as in a handshake.

Although the cytoplasmic domain of Tom20 is visible at a lower contour level in all four volumes, the resolution in those regions was not sufficient for building an atomic model. Nevertheless, we confirm that Tom20 adopts four conformations: two central, one bent and one extended (**Figure 4A**). The two central positions match one of the *N. crassa* Tom20 positions when superposed to the core of the complex (EMDB: 15850) **(Fig. S16)** (14). The bent conformation shows Tom20 interacting with Tom5 of the opposing protomer and matches the other observed *N. crassa* Tom20 conformation closely (EMDB: 15856) **(Fig. S16)** (14). In the extended conformation Tom20 opens towards Tom5 within the same protomer. As noted before (14), Tom20 appears to be highly dynamic (**Figure 4B**), interacting with Tom5 of both protomers, which has been suggested to be important in the transfer of preproteins from Tom20-22 to Tom40 (29). In contrast, in the resting conformation, both Tom20 receptors extend towards the center and appear to be immobile (**Figure 4C**). We note that the opposite Tom20 appears static in this scheme (**Figure 4B**). However, as the particles were preferentially aligned to one protomer by the cryoEM software, it is likely that both Tom20s move.

## Discussion

The structural changes underlying the protein translocation mechanism through TOM remain unclear despite the availability of several cryoEM structures at varying resolution (9–16). Here, we were able to determine the structures of both the TOM core and holo complexes from isolated mitochondria of the thermostable fungus *C. thermophilum*. In contrast, TOM structures from yeast and humans indicate that the Tom20 receptor subunit is destabilized during purification from mitochondria (9–12, 15, 16). Prior to our work on *C. thermophilum*, we purified and solved a native TOM holo complex from *N. crassa* outer mitochondrial membrane vesicles (OMVs) (14). In this previous structure, the peripheral Tom20 subunit appeared to be highly dynamic and present in sub-stoichiometric amounts relative to the other subunits, which made image processing difficult. Nevertheless, native mass spectrometry revealed the presence of stable subcomplexes containing two copies of Tom20 (14).

Our current structure of the TOM core complex is the most highly resolved so far, as evident from the clear side-chain features and lipids. A comparison of the lipid positions in our structure with those in *N. crassa* (PDB: 8B4I) and *H. sapiens* (PDB: 7CP9) indicated that four of them are conserved (**Fig. S8**) (9, 14). Consistent with other published structures (9, 14), the central lipid in our map sits on the twofold axis of the dimeric complex and bridges both protomers. Within the yeast structure, which was of a complex prepared with a slightly harsher detergent, two dodecyl-β-D-maltoside (DDM) molecules had displaced the central lipid in a structurally analogous manner, with their sugar moieties stacked and their aliphatic tails extending along Tom22 and β1, β18 and β19 of Tom40 (10). By contrast, the central lipid is positioned further down towards the IMS along the twofold axis *in H. sapien*s and *D. melanogaster* (9, 13). The shift in lipid position may result from the markedly bent conformation of Tom22 (9, 13). We found additional lipids wedged between Tom6 and Tom7 that would strengthen the interaction of these small subunits with Tom40 (**Figure 1D-E**). We note that the head groups of the lipids were not entirely visible, suggesting that the hydrophobic acyl chains interact more strongly with the complex. We opted to model the TOM lipids as PC, as this bilayer-forming lipid is the most abundant in the OMM of *N. crassa* (30), followed by the non-bilayer-forming phosphatidyl ethanolamine (PE). Both lipids play different but equally important roles, with PE being critical for TOM function while PC is important for its biogenesis (31, 32). Although not visible in our structure, cardiolipin may be important for the association of Tom20 with the TOM core complex (33).

The parallel closure of strands β1 and β19 in the Tom40 β-barrel gives rise to a strained hydrogen-bond network. A weak seam of the β-barrel has been proposed to be a mechano-sensitive feature of TOM that may play a role in preprotein translocation (34). However, we find no evidence in the structure that Tom40 can undergo such drastic changes. The barrel seam would be stabilized by the interaction with Tom22 and the central lipid, making large-scale conformational changes energetically costly and hence unlikely. The observation of potential water clusters between the inner helix and the inner barrel wall supports our previous findings that this region is comparatively rigid (**Fig. S9**) (35), consistent with the highly conserved Tom40 residues in this position (36).

At 3.2 Å resolution, our map of the TOM holo complex shows how Tom20 docks to the native core complex. Compared to the previous structure of *N. crassa* TOM, with one Tom20 resolved in the dimer, our current *C. thermophilum* structure indicates two symmetrically bound Tom20s. The docking position of Tom20 in relation to Tom22 at the cytoplasmic-membrane surface between the two species is unchanged. Through comparative sequence analysis, we demonstrated that Tom20-Tom22 exhibits a complementary electrostatic interaction that is conserved across fungi and metazoa **(Fig. S11)**. The transmembrane helices of *C. thermophilum and N. crassa* Tom20 are flexible and, therefore, not well resolved. We speculate that this helix flexibility is a necessary feature of the subunit, whereby it counters the movement of the receptor domain in the cytoplasm, enabling it to translocate preproteins efficiently. Given that the Tom20 transmembrane helix lacks direct contact with the rest of the core complex, we propose that in the OMM, it is stabilized solely by lipid interactions. Lipids are crucial for TOM transport activity and biogenesis (31, 32). In particular, cardiolipin has been shown to be critical, in that cardiolipin-deficient yeast mutants display impaired assembly of Tom20 with the rest of the complex (33). We propose that the stability of the attachment of Tom20 is a result of the thermostable nature of *C. thermophilum*.

In the crosslinked human TOM holo complex (15), Tom20 looks strikingly different from the un-crosslinked *N. crassa* and *C. thermophilum* structures **(Fig. S15, S17A-B)** and from Tom20 in the recent native *H. sapiens* TOM-PINK1-VDAC complex (17) (PDB: 9EIH) **(Fig. S17C)**. In particular, the human crosslinked structure shows a single direct interaction between the transmembrane helices of Tom22 and Tom20. In our structure, this presumed Tom20-Tom22 contact site coincides with lipid binding sites that appear to be highly conserved. Bound lipids in this position would prevent such an interaction. Furthermore, *in vivo* photo-crosslinking and targeted protease treatment in yeast has shown clearly that the Tom22 receptor domain is primarily required for integrating Tom20 into the complex (37, 38). The recent structure of the human TOM-PINK1-VDAC (17) contains a single Tom20 copy per complex that agrees with the position of Tom20 in our structure **(Fig. S17B)**. By binding to the cytoplasmic domain of Tom22 instead of its transmembrane helix, Tom20 can be flexible and function synergistically with Tom22 in presequence recognition and translocation.

On the cytoplasmic side, the TPR domain of Tom20 closely resembles that of the published crystal structure of rat Tom20 (26). As in our map of the TOM complex, the crystal structure reveals a bound pALDH presequence, with its amphipathic helix nestled within the hydrophobic patch on the receptor surface. Consistent with this observation, NMR studies (26) have shown that presequences equilibrate between different binding states, which might enable Tom20 to recognize diverse presequences. In our structure, two receptor domains are poised at the center of the complex, with the hydrophobic segments of their TPRs orientated toward each other. We refer to this as the ‘handshake’ conformation (see **Fig. 4C**), in which both Tom20s rest at the center of the complex. We speculate that in this handshake conformation, both Tom20 receptors interact with each other, awaiting the arrival of the incoming preprotein, whereupon they engage in translocation together with Tom22 (37). Furthermore, we note that the positions of helices α2-6 in our Tom20 handshake conformation differ from those in the human TOM-PINK1-VDAC array, as Tom20 interacts with PINK1 (17). This suggests that the TOM complex and its subunits serve a different role in mitochondria beyond preprotein translocation (**Fig. S17**).

Within the translocation pore, we identified several densities of the presequence of pALDH, consistent with the *N. crassa* structure (14). The lower-resolution densities match the presequence binding sites along the acidic patch of Tom40 identified by photo crosslinking (28), indicating that substrate binding along this pathway is conserved. The multiple copies of bound pALDH in our structure may suggest that each pore can simultaneously translocate several preproteins, even though this might result in unfavorable molecular crowding. Regarding the location and structure of Tom70 in the complex, more work remains to be done.

Finally, we find that, in addition to the resting handshake conformation, Tom20 can adopt four distinct conformations on the cytoplasmic membrane surface. Two of these states match those we see in *N. crassa* TOM (14). One particular state of Tom20 extends to Tom5 and the longest loop of Tom40, both of which have been implicated in protein translocation (14, 29). Tom20 is an essential receptor in mitochondrial protein translocation. Many functionally and structurally analogous receptors appear independently in eukaryotic lineages, apparently as a result of convergent evolution (39). For example, the equivalent but unrelated plant Tom20 and the atypical TOM46 (ATOM46) of *Trypansoma brucei* both have cytoplasmic TPR and armadillo repeat receptor domains (40, 41) and work as protein-binding modules.

In conclusion, our work highlights the molecular basis of Tom20 interaction with the TOM core complex through Tom22. The configurations of the Tom20 receptor we observe are likely to respectively represent an idle state and different dynamic states in the early steps of preprotein translocation. We find that our presequence adopts various binding modes in the twin-translocating pore and in the Tom20 receptor. Promiscuous preprotein binding provides us with insights into how TOM functions as the central protein transport module into submitochondrial compartments.

## Materials and Methods

### Generation of a genetically modified *C. thermophilum* strain

To enable the purification of the TOM complex from *C. thermophilum*, we first synthesized (GenScript) a C-terminal FLAG-tagged variant of the gene encoding Tom22 (CTHT_0026640). The synthesized gene was cloned together with a 750 bp upstream Actin (CTHT_0062070) promoter and a 300 bp downstream *GAPDH* (CTHT_0004880) terminator region into a pNK51 vector (with the *ERG1* thermostable selection marker), which was kindly provided by Dr. Nikola Kellner and Prof. Ed Hurt (Biochemistry Center, Heidelberg University, Germany). The resulting plasmid was linearized by restriction enzyme digest, concentrated through ethanol precipitation and stored at -20 °C until further use.

*C. thermophilum* was transformed with the linearized plasmid using the method developed by Kellner *et al* (42). Briefly, wild-type *C. thermophilum* spores (DSM 1495) were initially re-germinated on a CCM agar plate at 52°C for 2-3 days. Mycelia were scraped from the plate and small pieces used to inoculate CCM media for scaling up to a 150 ml culture. Mycelia were then harvested and digested using an enzyme blend comprising pectinases, beta-glucanase, protease, and arabinanase (Vinotaste Pro, Novozymes). The protoplasts were filtered from the digested mycelia and subsequently used for PEG-mediated transformation with 10 μg of the linearized plasmid DNA. The transformed protoplasts were instantly plated on a double-layer CCM+Sorbitol (CCMS) plate, comprising a top layer of CCMS agar and a bottom layer of CCMS agar supplemented with 0.5 μg/ml terbinafine, and grown at 50 °C for 5-7 days. Surviving transformants were further selected by transferring to a new CCM agar plate supplemented with 0.5 μg/ml terbinafine to confirm stable gene integration. Stable transformants were utilized for subsequent small-scale culture, and the target protein was detected by western blotting with the monoclonal anti-FLAG M2 antibody (Sigma-Aldrich, F3165). Spores were generated from successful transformants for long-term storage at -20 °C as described (42).

### Large-scale growth of *C. thermophilum* Tom22 strain

Spores for *C. thermophilum* expressing Tom22-FLAG fused with a FLAG epitope were re-germinated on a CCM agar plate containing 0.5 μg/ml terbinafine for 3 days at 52 °C. Mycelia were scraped from the plate, finely chopped and grown in 400 ml CCM media supplemented with 100 µg/ml of ampicillin for 24 hr at 85 RPM and 52°C in a rotary incubator. The resulting pre-culture was blended using a GrindoMix GM300 (Retsch) at 4,000 RPM for 2x 2 min. 6x 2 l CCM in 5 l baffled flasks were individually inoculated with 100 ml blended pre-culture and incubated for 15-18 hr at 75 RPM and 52 °C. To harvest the cultures, *C. thermophilum* mycelia were collected in a sieve, dried and frozen as granules in liquid nitrogen. The granules were stored at -70°C until further use. Typically, one preparation yielded 22 g of dried granules.

### Preparation of mitochondria

Mitochondria were isolated as previously described for complex I (43). In brief, thawed *Chaetomium* cell pellets were resuspended at a 9:1 (w/v) ratio of cells to solution using *Ct* mitochondrial buffer composed of 50 mM HEPES (VWR), 350 mM sorbitol (VWR), 1 mM EGTA (Sigma) and 1 mM PMSF (Sigma). To fractionate the resuspension, cells were sonicated on ice using a Branson 250D Sonifier (G.Heinemann Ultraschall und Labortechnik) 2x 2 min (3 s and 2 s pulse on and off, respectively) with a 5 mm tip and set to a 25-30% amplitude with a 2 min rest period. The homogenate was centrifuged at 1,500 x*g* for 5 min and the supernatant was decanted. The supernatant was further clarified by centrifugation at 4,000 x*g* for 5 min. Mitochondria were pelleted at 12,000 x*g* for 15 min and resuspended in 30 ml SME buffer. The step was repeated to increase purity. Mitochondria were diluted to 5-10 mg/ml and snap-frozen in liquid nitrogen. Mitochondria were stored at -70°C until further use. Each preparation yielded 30 mg to 46 mg of mitochondria for protein purification.

### Purification of substrate-free TOM complex

The TOM complex was purified from isolated mitochondria according to the protocol established for *N. crassa* (*14, 44*). First, mitochondrial protein complexes were extracted at 1 mg/ml in solubilization buffer containing 20%(v/v) glycerol, 10mM MOPS (VWR) pH 7.0, 50 mM KAc (VWR), 50 mM imidazole (Sigma), 1 mM PMSF and 1% GDN (w/v, Anatrace). The sample was then left to incubate under rotation for 1 h at 4°C. The lysate was subjected to centrifugation at 13,000 x g for 10 min. The clarified supernatant was mixed with anti-FLAG M2 affinity gel (Sigma) and incubated overnight under rotation at 4°C. To wash off unbound contaminants, the resin was washed in a gravity flow column (BioRad) using 15 ml of 10 mM MOPS pH 7.0, 50 mM potassium acetate, 1 mM PMSF and 0.02% GDN. The TOM complex was eluted in three 500 µl fractions using the same buffer, with the addition of 0.45 mg/ml 3XFLAG peptide (Sigma-Aldrich). The three eluted fractions were pooled and concentrated to 100 µl using an AmiconUltra 100 kDa MWCO (Millipore). The TOM complex was further purified by injecting the concentrated sample onto a Superdex 200 Increase 5/150 GL (Cytiva) and eluted at a flow rate of 0.1 mL/min in size-exclusion buffer: 50 mM potassium phosphate pH 8.0, 50 mM KCl, 1 mM EDTA, 1 mM TCEP and 0.02% GDN. TOM eluted at 1.1 ml at a peak concentration of 1.64 mg/ml. The sample was immediately used for cryoEM grid preparation.

### Purification of pALDH-bound TOM complex

To obtain a structure of the pALDH-bound complex, purified TOM was incubated with a synthesized mitochondrial targeting sequence of pALDH (GenScript), as described (14). In brief, isolated TOM and pALDH were mixed at a ratio of 1: 8, and left for 1 h. Unbound pALDH was removed by size exclusion chromatography using a Superdex 200 Increase 5/150 GL column in size exclusion buffer at a flow rate of 0.1 ml/min. Peak fractions containing TOM were pooled and concentrated (AmiconUltra 100 kDa cutoff) to 25 µl at 1.9 mg/ml. The purified complex was used immediately for cryoEM grid preparation.

### CryoEM preparation and imaging

Grids were plunge-frozen using a Vitrobot Mark IV (Thermo Scientific) operating at 4°C, 100% relative humidity and a blot force of 20. Roughly 3 µl of substrate-free or pALDH-bound samples were applied to glow-discharged holey carbon R2/2 Cu grids (Quantifoil) and blotted for 5-7 or 3 s, respectively. The grids were then plunged into liquid ethane.

For TOM without bound preprotein, 15,566 movies were collected on a Titan Krios G3i (Thermo Scientific) transmission electron microscope operating at 300 keV and equipped with a BioQuantum-K3 imaging filter (Gatan), using an energy slit width of 20 eV. Dose-fractionated movies were collected at a pixel size of 0.837 Å corresponding to 105,000x nominal magnification. The total accumulated dose was 60 e^-^/Å^2^ over 60 frames after an exposure time of 3.3 s. The defocus range was set from -0.9 to -2.4 µm.

The pALDH-bound complex was imaged using a Krios G4 transmission electron microscope operating at 300 keV and equipped with a E-CFEG cold field emission gun, a Selectris X energy filter, and a Falcon 4i camera (Thermo Scientific). 34,438 movies were acquired in EER format at a pixel size of 0.573 Å corresponding to 215,000x nominal magnification. The total accumulated dose was 70 e-/Å^2^ after an exposure time of 3 s. The defocus range was set from -1.6 to -2.4 µm.

### Image processing

To account and correct for beam-induced motion and radiation damage, the Relion 3.0 implementation MotionCorr was applied to 15,566 movies of the substrate-free TOM complex (45). Micrographs were manually curated and 14,334 retained. Curated micrographs were then imported into CryoSPARC v4.0 (46), where they were subjected to patch CTF correction. A combination of particle selection algorithms was used on the micrographs including crYOLO (47), Topaz (48), CryoSPARC v4.0 blob picker and template matching. The latter three were used within CryoSPARC v4.0 (46), while crYOLO was used as a standalone application software. All further processing steps for both datasets was carried out in CryoSPARC v4.0 (46). In parallel, particles picked with different software packages were first extracted and downsampled to a pixel size of 3.6 Å with a box size of 84 pixels. To remove duplicates or particles unsuitable for high-resolution cryoEM, iterative rounds of 2D classification resulted in 1,379,719 picks. Particles were re-extracted and downsampled to 1.7 Å pixel size and a box size of 180 pixels. Then, a further round of 2D classification and *ab initio* reconstruction was carried out, resulting in 478,043 particles. To resolve different conformational states of the TOM complex, particles were re-extracted without downsampling with a box size of 360 pixels, then separated into ten classes by *ab initio* reconstruction followed by heterogeneous refinement. Five classes with high-resolution reconstruction of TOM, together consisted of 326,681 particles, were combined and further processed using non-uniform refinement with imposed C2 symmetry, resulting in a structure with a global resolution at 3.2 Å based on the gold-standard FSC 0.143 criterion (49). The TOM core complex was resolved to 3.17 Å using an appropriate mask and by applying local refinement to the particles, with C2 symmetry imposed. The TOM holo complex was resolved to 3.75 Å by applying non-uniform refinement with imposed C2 symmetry to a single class of 76,100 particles from the previous heterogeneous refinement.

Images of the preprotein-bound dataset were processed using CryoSPARC v4.0 (46). Following patch motion correction and CTF estimation, the dataset was manually curated to 31,468 micrographs. Particles were initially blob-picked and used to train a Topaz model (48), resulting in 1,443,703 extracted particles. The dataset was then classified in 2D to discard artifacts. The best 826,204 particles were further separated into four *ab-initio* reconstructions and heterogeneously refined without imposed symmetry. The best class, containing 345,380 particles, was non-uniformly refined without imposed symmetry, resulting in a 2.93 Å consensus map **(Fig. S2)**. A map of the TOM core complex was obtained after subsequent refinements of this dataset. A final local refinement with imposed C2 symmetry resulted in a 2.67 Å resolution map, as assessed by the gold-standard FSC 0.143 criterion **(Fig. S5)**. The consensus particles were further separated into ten clusters via 3D variability analysis (50), without imposed symmetry, by masking only the cytoplasmic side of the complex. Particles belonging to two clusters were pooled. Duplicates were removed, amounting to 51,299 particles **(Fig. S2)**. These were further non-uniformly refined with imposed C2 symmetry, resulting in a 3.15 Å resolution map of the TOM holo complex, as assessed by the gold-standard FSC 0.143 criterion **(Fig. S10)**. A parallel non-uniform refinement of the same particles under symmetry relaxation revealed the presence of pALDH bound to the TOM holo complex. The preprotein density was isolated by subtraction of the unbound TOM holo map from the bound map using UCSF ChimeraX **(Fig. S14)** (51).

### Model building

To generate atomic coordinates for the TOM core complex, *de novo* model building was carried out using ModelAngelo (52) with the 2.67 Å resolution TOM core consensus map and corresponding amino acid sequences serving as input. Water and lipid molecules were added in Coot where appropriate, as defined by clear density and chemical coordination (53). The model was then refined against the substrate-free and substrate-bound TOM core maps using ISOLDE (54). The TOM holo model was generated using a combination of the previous core model and AlphaFold3 (25) to generate coordinates for the Tom20 subunit. UCSF ChimeraX was used to dock the subunits in the substrate-free and substrate-bound holo complex before refinement in ISOLDE (51, 54). Finally, all models were globally refined in Phenix with global minimization and B-factor (ADP) refinement (55).

## Acknowledgments

We acknowledge Nikola Kellner and Ed Hurt for the pNK51 vector. We thank Janet Vonck for advice on modeling. We thank Juan Castillo and Özkan Yildiz for computational support, as well as Sonja Welsch and the team of the Central Electron Microscopy Facility of the Max Planck Institute of Biophysics for cryoEM support.

## Data and materials availability

The cryoEM maps and models were deposited under the following accession code: preprotein-bound TOM core (PDB: 9I6B, EMD-52652), preprotein-bound TOM holo (PDB: 9I7S, EMD-52660), preprotein-free TOM core (PDB: 9I7P, EMD-52658) and preprotein-free TOM holo (PDB: 9I7T, EMD-52661).

---

**Figure S1.**
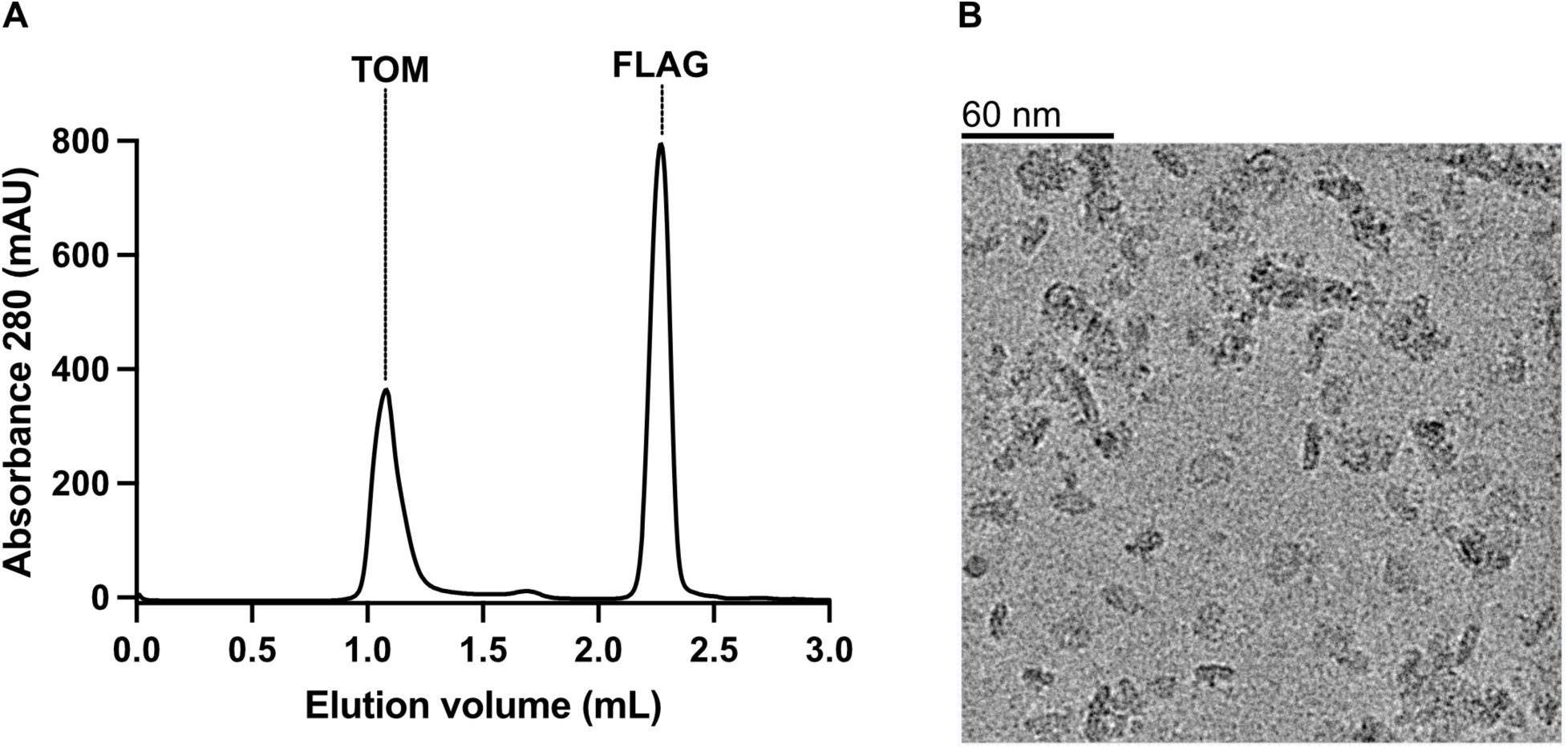
Purification of *Chaetomium thermophilum* TOM from isolated mitochondria. (**A**) Size-exclusion chromatography profile using a Superdex 200 Increase 5/150 GL for the purified TOM complex separated from the FLAG peptides. (**B**) An exemplary cryoEM micrograph with the scale bar above the image.

**Figure S2.**
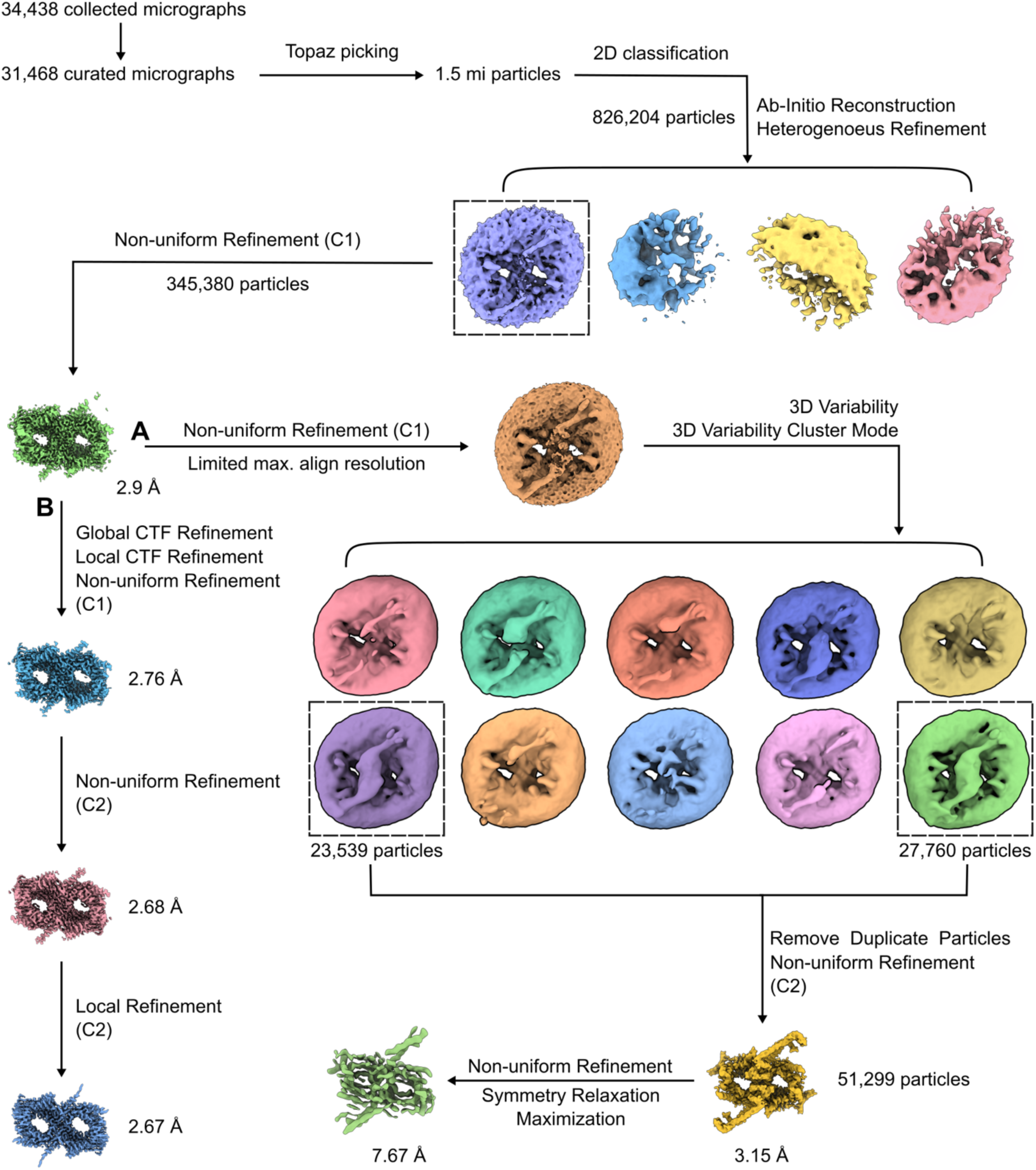
CryoEM processing pipeline for the pALDH bound dataset. (**A**) The workflow that resulted in two Tom20 bound C2-symmetrised and symmetry-relaxed holo complexes at resolutions of 3.15Å and 7.67Å, respectively. (**B**) The processing pathway that yielded the TOM core structure at 2.67Å resolution.

**Figure S3.**
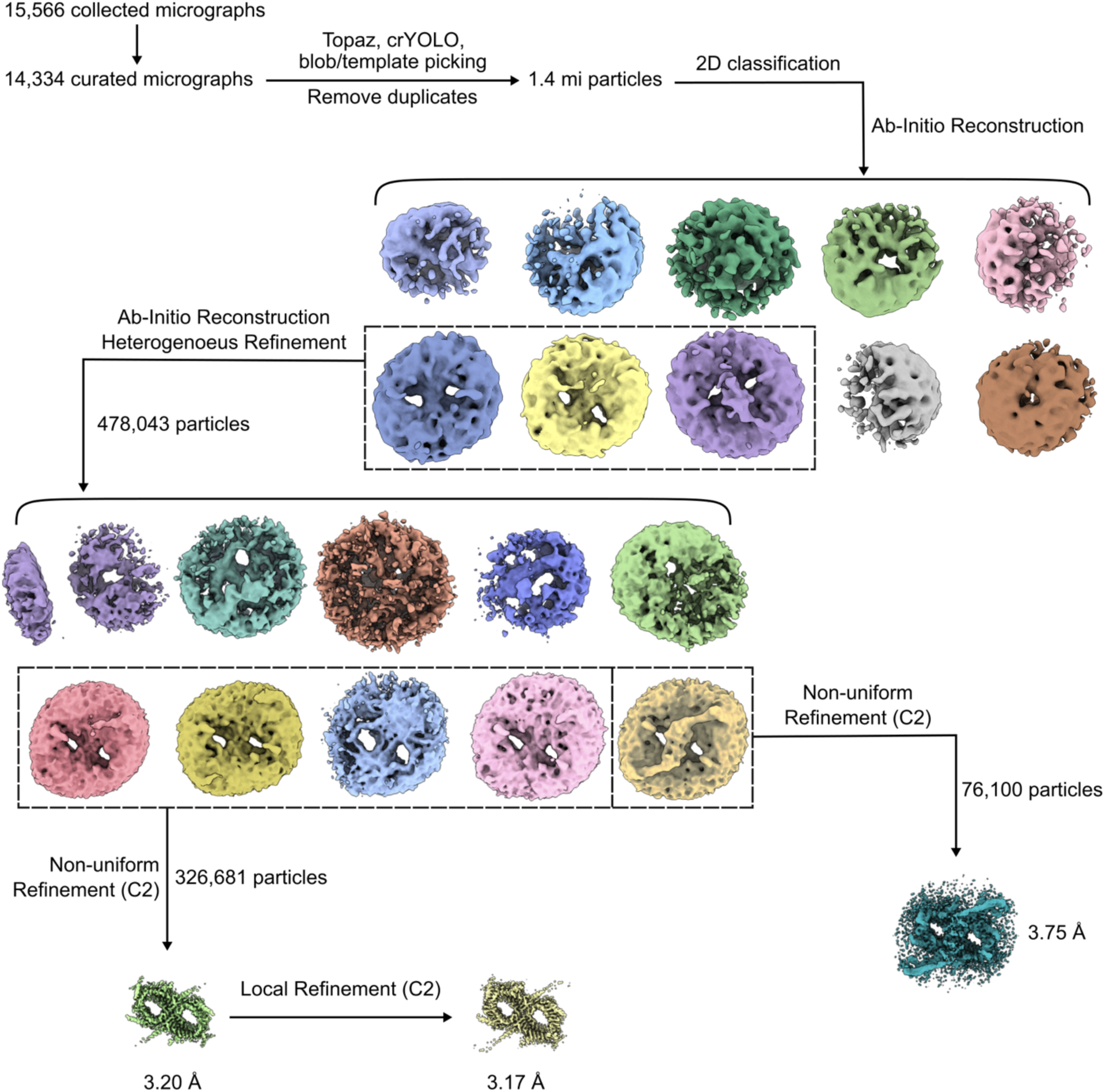
CryoEM processing pipeline for the substrate-free dataset. The workflow yielded the C2-symmetrized TOM core and holo complexes at 3.17Å and 3.75Å resolution, respectively.

**Figure S4.**
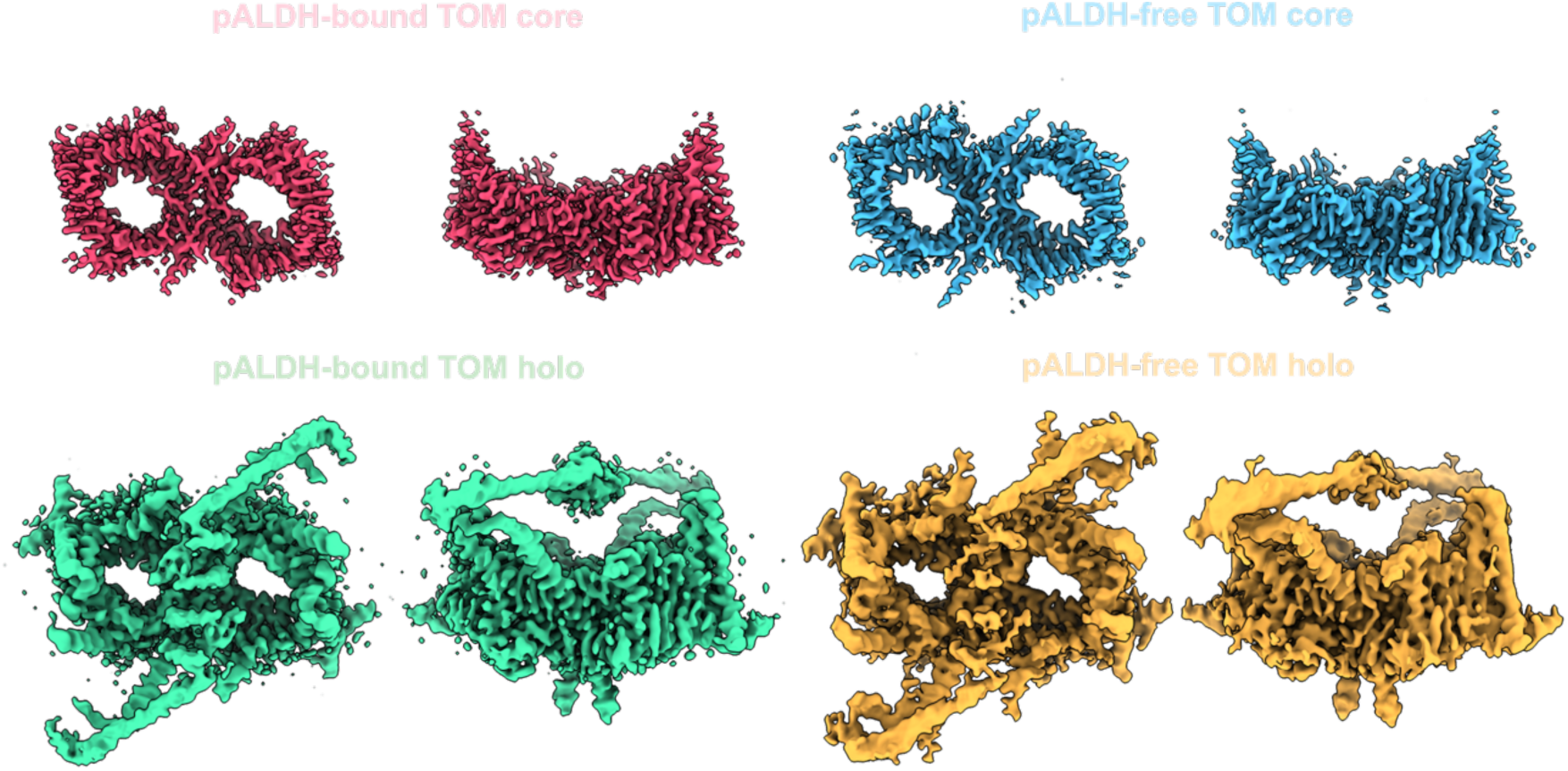
Comparisons of the final cryoEM reconstructions of the *C. thermophilum* TOM complexes. The volumes were viewed in UCSF ChimeraX (1), and the ‘hide dust’ function was applied only to pALDH-free TOM with a size limit set to 5 Å.

**Figure S5.**
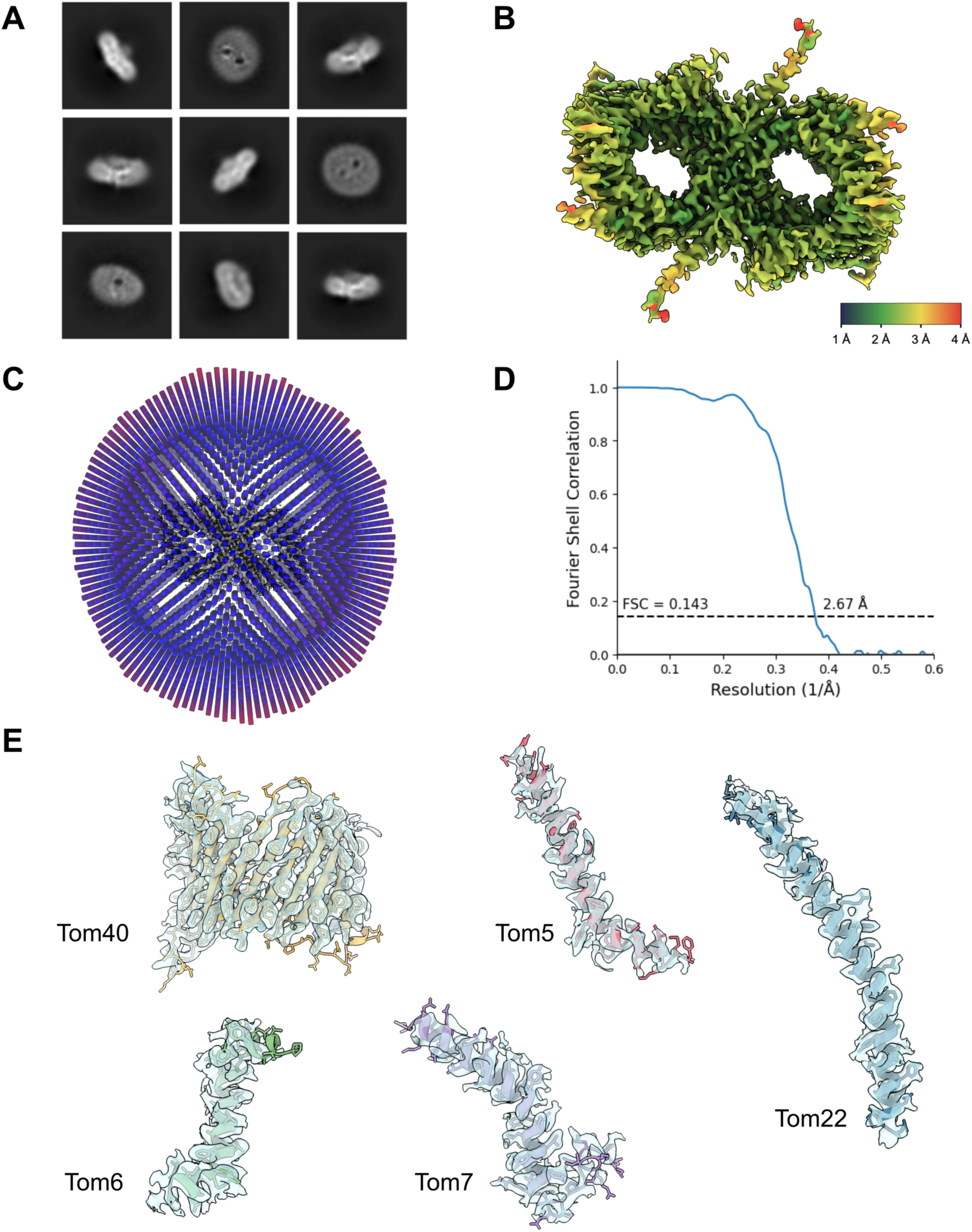
Quality assessment of the *C. thermophilum* TOM core single-particle dataset. (**A**) 2D class averages of the TOM holo complex. (**B**) Local resolution estimation of the reconstruction with a scale bar below to indicate resolution values. **C**) The angular distribution of the particles in the dataset is displayed over the volume with the length of the bar indicating abundance. (**D**) The corrected FSC curve of the reconstruction with the resolution was determined at a threshold of 0.143. (**E**) The cartoon representation of the subunits of the TOM core complex with their cryoEM densities.

**Figure S6.**
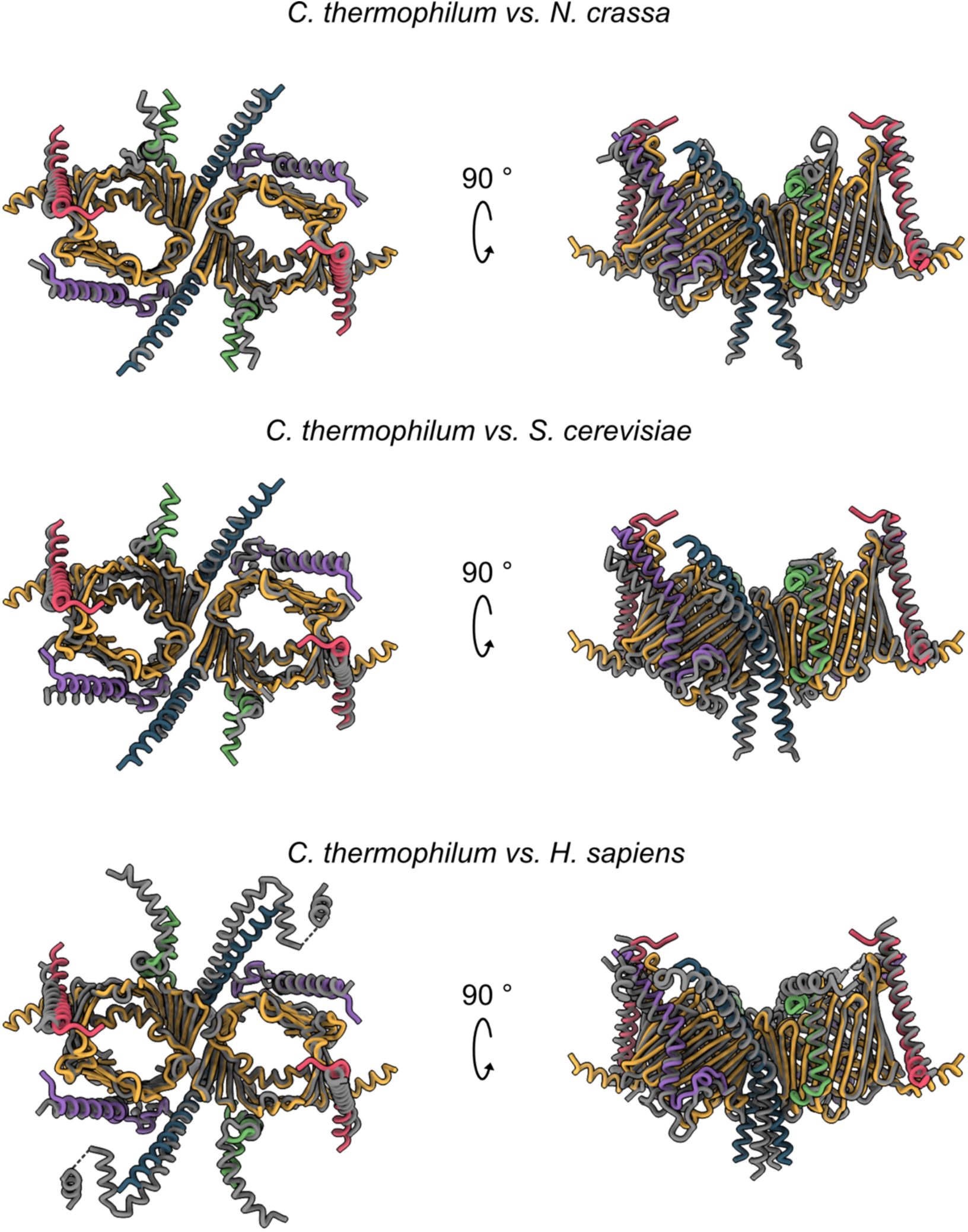
Structural comparison of the *C. thermophilum* TOM core complex with other species. Structures from other species (gray) were superposed onto that of *C. thermophilum* (colored) in UCSF ChimeraX and displayed as cartoons. The *C. thermophilum* subunits are colored yellow (Tom40), blue (Tom22), purple (Tom7), green (Tom6) and pink (Tom5). The RMSD values between *C. thermophilum* and *N. crassa* (8B4I), *S. cerevisiae* (6UCU), and *H. sapiens* (7CP9) were 0.725 Å (based on 286 atoms), 0.812 Å (based on 268 atoms), and 0.840 Å (based on 214 atoms).

**Figure S7.**
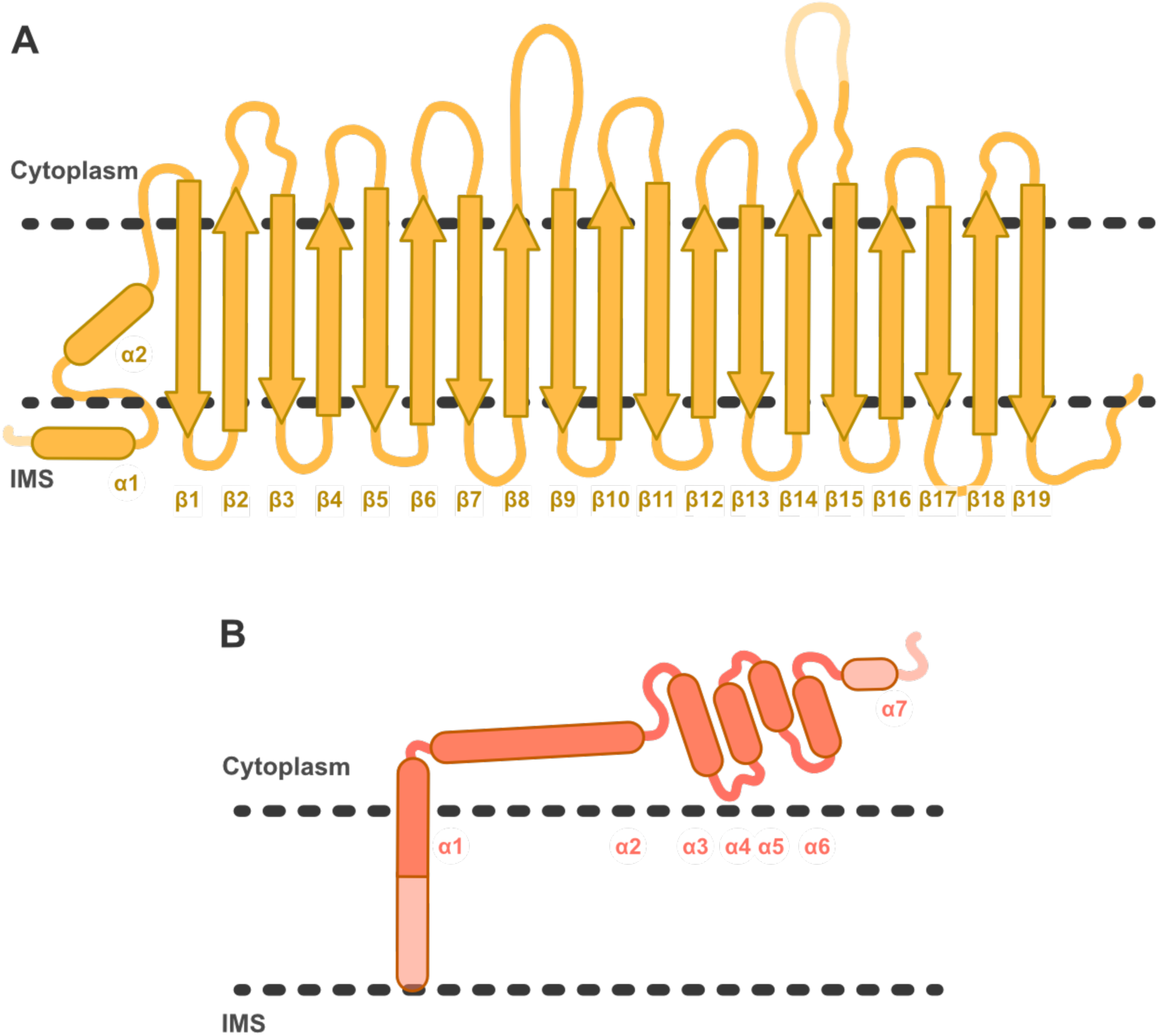
The topology of the *C. thermophilum* Tom40 and Tom20 subunits. (**A**) A schematic representation of the nineteen β-strands, the interconnecting loops and the three α-helices. (**B**) A schematic representation of the six α-helices of Tom20, including the tetratricopeptide repeat (TPR) fold α3-6. Unresolved regions are shown in a lighter color.

**Figure S8.**
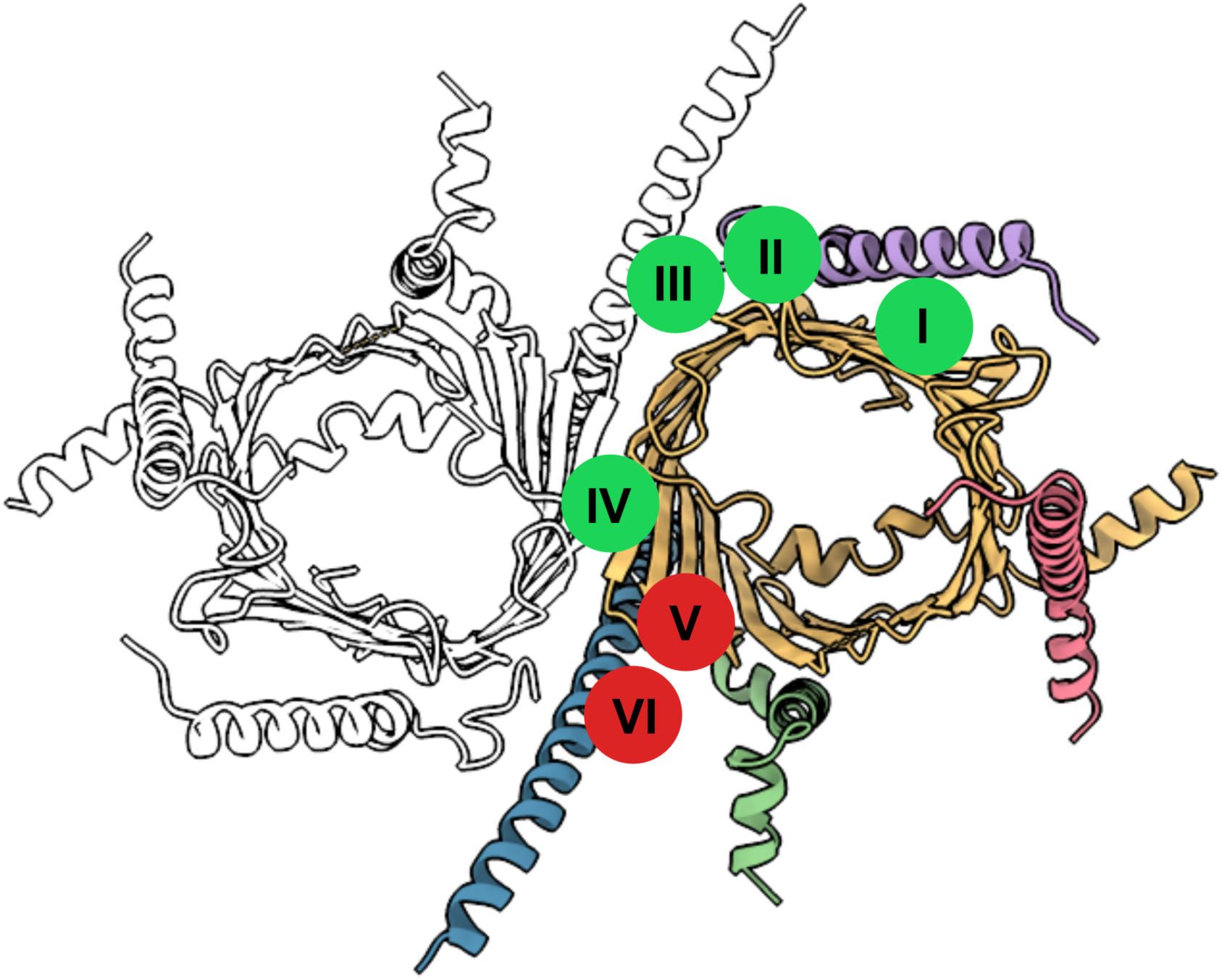
Conservation of lipid positions across organisms. The *C. thermophilum* TOM complex is represented in cartoon with only one protomer colored. Lipid positions in *C. thermophilum* (numbered) are compared with those of *N. crassa* (PDB: 8B4I) and humans (PDB: 7CP9). Green and red circles indicate whether those positions are conserved or not, respectively.

**Figure S9.**
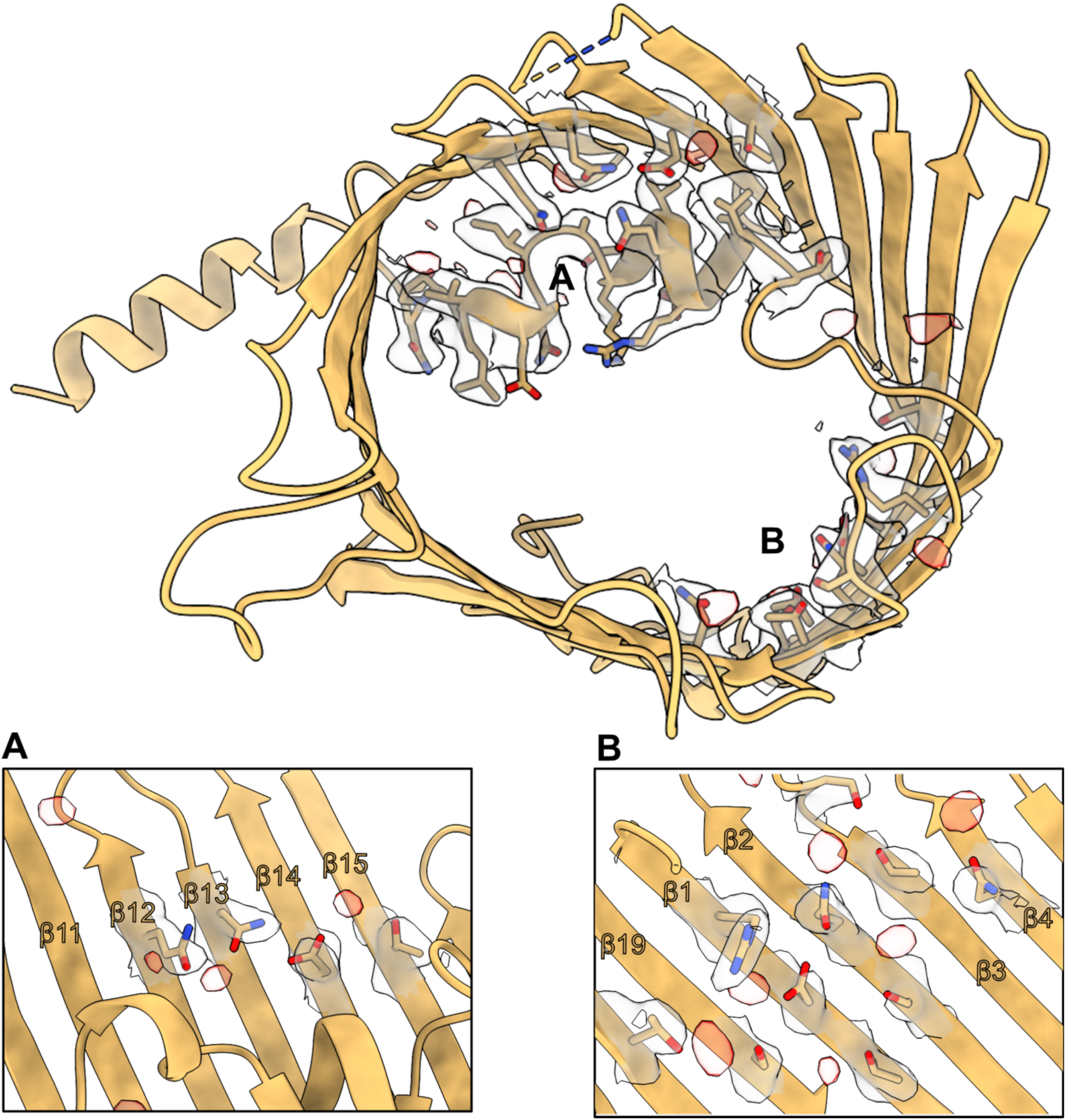
Clusters of unassigned waters or ions. (**Top**) Densities of suspected waters or ions (red) observed within the Tom40 pore from the cytoplasmic side. (**A**-**B**) Close-up of amino acid residues adjacent to the unassigned densities.

**Figure S10.**
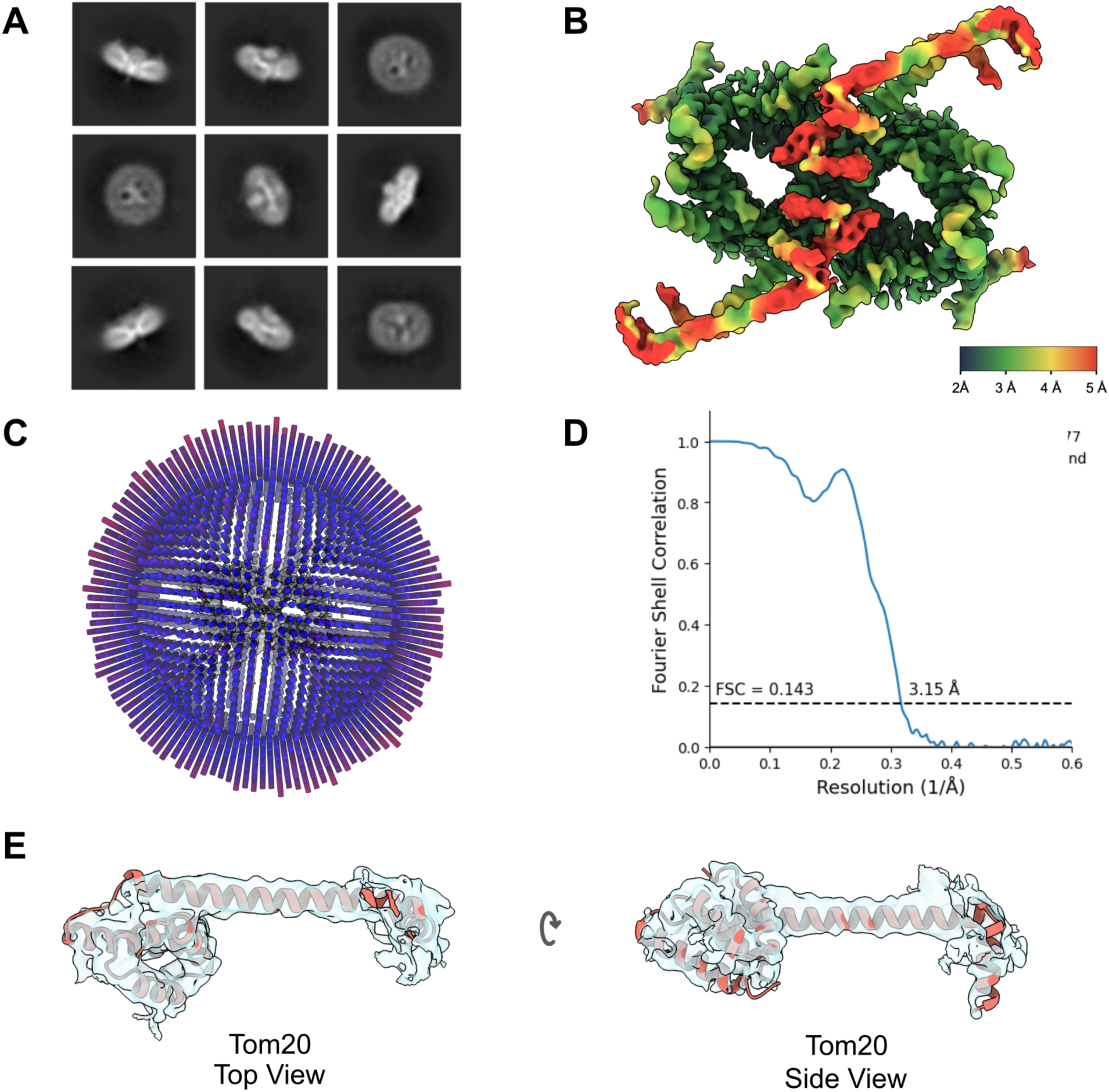
Quality assessment of the *C. thermophilum* TOM holo single-particle dataset. (**A**) 2D class averages of the TOM holo complex. (**B**) Local resolution estimation of the reconstruction with a scale bar below to indicate resolution values. (**C**) The angular distribution of the particles in the dataset is displayed over the volume with the length of the bar indicating abundance. (**D**) The FSC curve of the reconstruction with the resolution was determined at a threshold of 0.143. (**E**) The cytoplasmic portion of the Tom20 subunit in cartoon representation with its cryoEM density.

**Figure S11.**
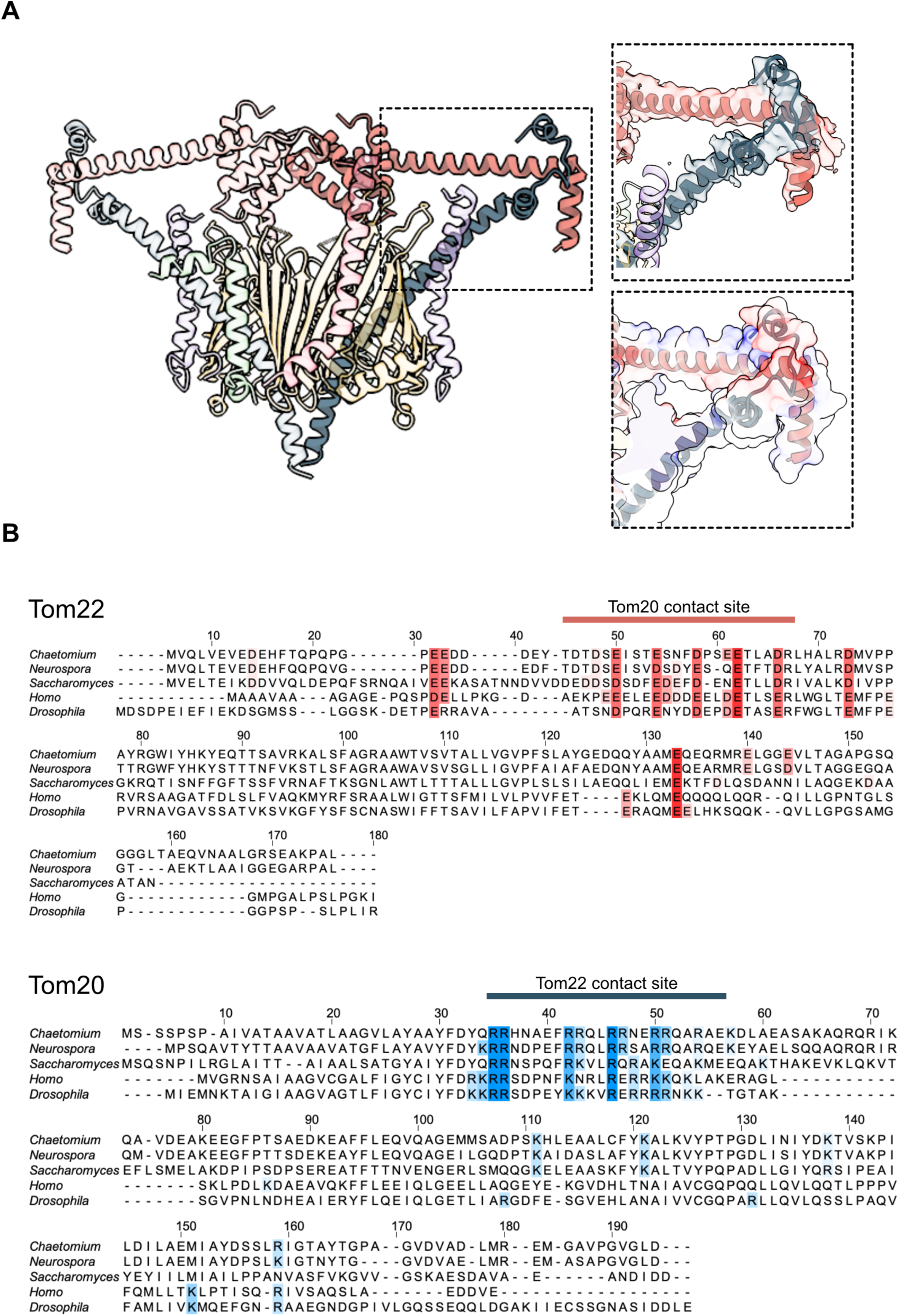
Molecular interactions of Tom20 with Tom22. (**A**) The TOM holo complex is depicted as a cartoon representation with one of the Tom22 (blue) and Tom20 (red) displayed as opaque. The dashed boxes highlight the interaction between the two subunits, with the close-up showing their corresponding cryoEM density (top) or the model-derived coulombic electrostatic potential (bottom). (**B**) Sequence alignments of Tom22 and Tom20 across various organisms. Conserved residues for the Tom22 and Tom20 contact sites are highlighted in red and blue, respectively.

**Figure S12.**
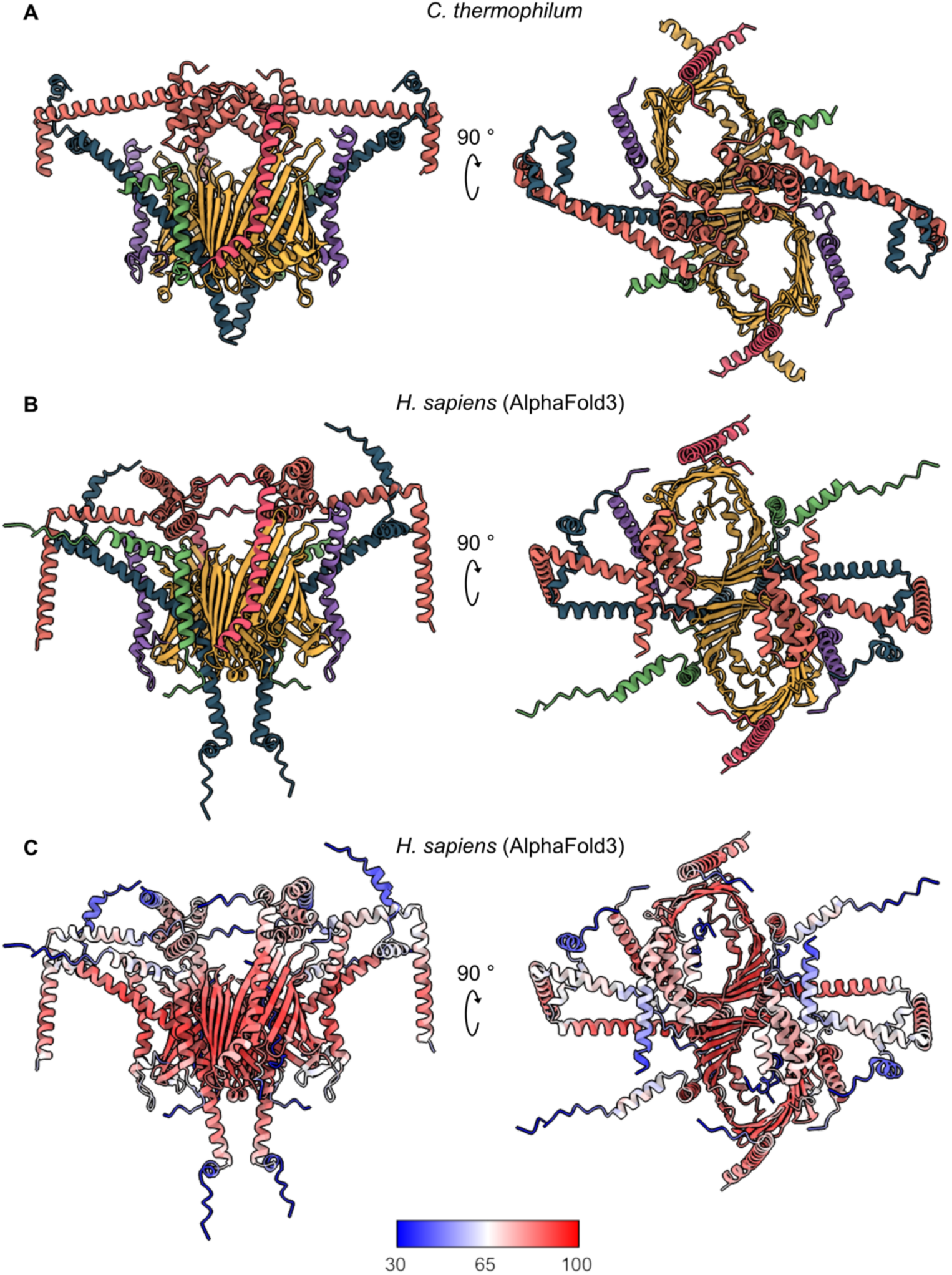
The symmetrical TOM holo complex is not unique to *C. thermophilum*. A visual comparison between the *C. thermophilum* (**A**) and the AlphaFold3 (2) predicted TOM holo complex from *H. sapiens* (**B**). (**C**) The predicted local distance difference test (pLDDT) is represented on the model with a corresponding scale bar to indicate its value.

**Figure S13.**
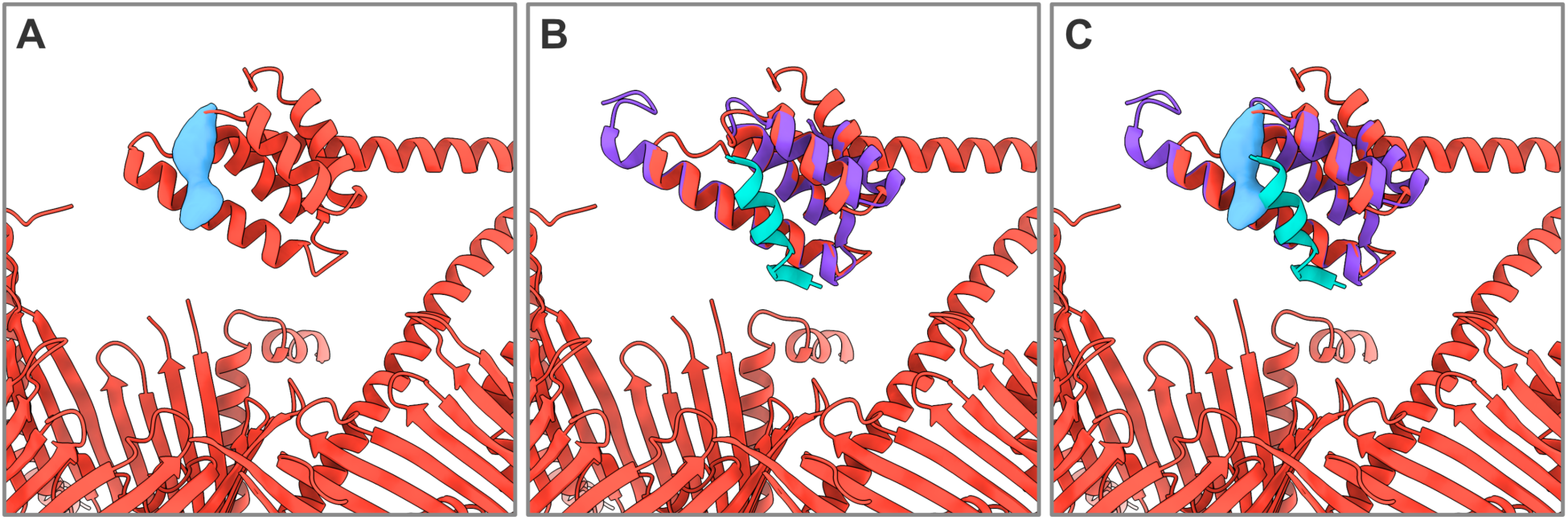
Tom20 TPR domain and receptor-preprotein interaction. (**A**) A bound preprotein density (blue) is shown bound to the *C. thermophilum* Tom20 structure (red). (**B**) The rat crystal structure of Tom20 (purple, PDB: 3AWR) with bound pALDH (cyan) is superposed to the *C. thermophilum* structure. (**C**) An overlay of (**A**) and (**B**).

**Figure S14.**
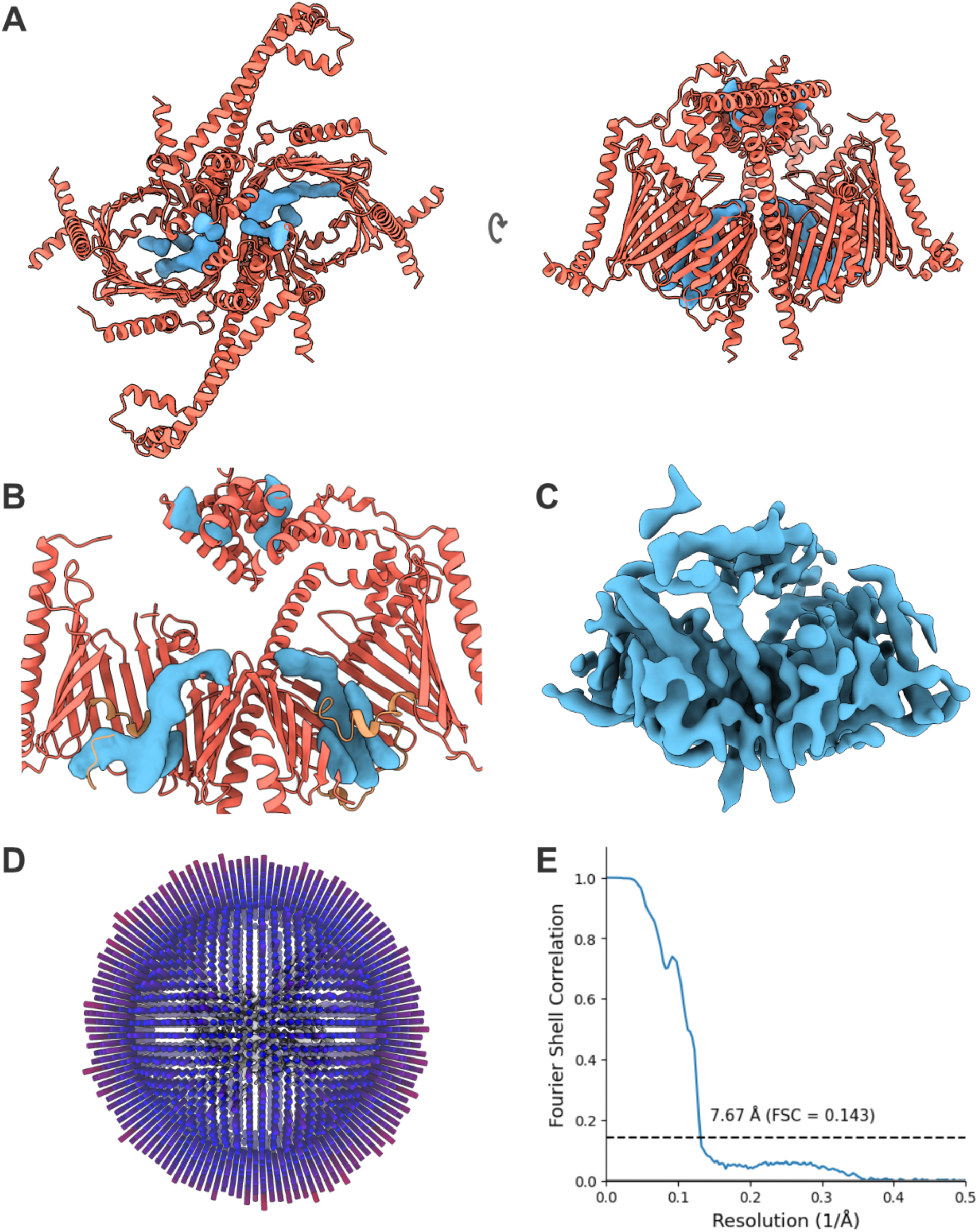
Preprotein binding to the TOM holo complex. (**A**) Preprotein densities displayed alongside the TOM holo model were generated by subtraction of the unbound TOM holo map from the bound map using UCSF ChimeraX. (**B**) A close-up and sectional view of preprotein densities. (**C**) Non-subtracted cryoEM density of the pALDH-bound TOM holo complex. (**D**) Angular distribution of particles from the symmetry relaxed dataset (**Figure S2**). (**E**) The FSC curve of the reconstruction where the resolution is determined at a threshold value of 0.143.

**Figure S15.**
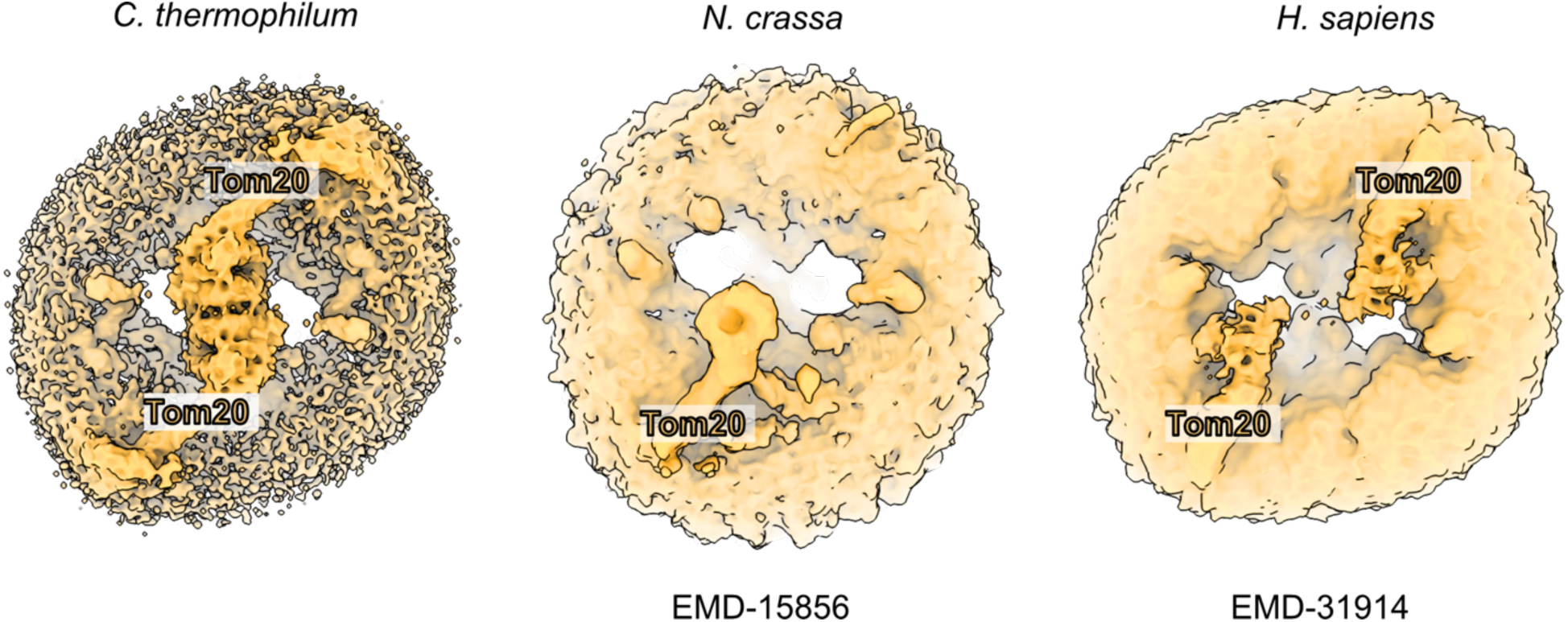
Comparison of cryoEM maps of the TOM holo complex across species. Contour levels for each cryoEM reconstruction were adjusted in UCSF ChimeraX (1) to ensure that Tom20 (labeled) would be visible.

**Figure S16.**
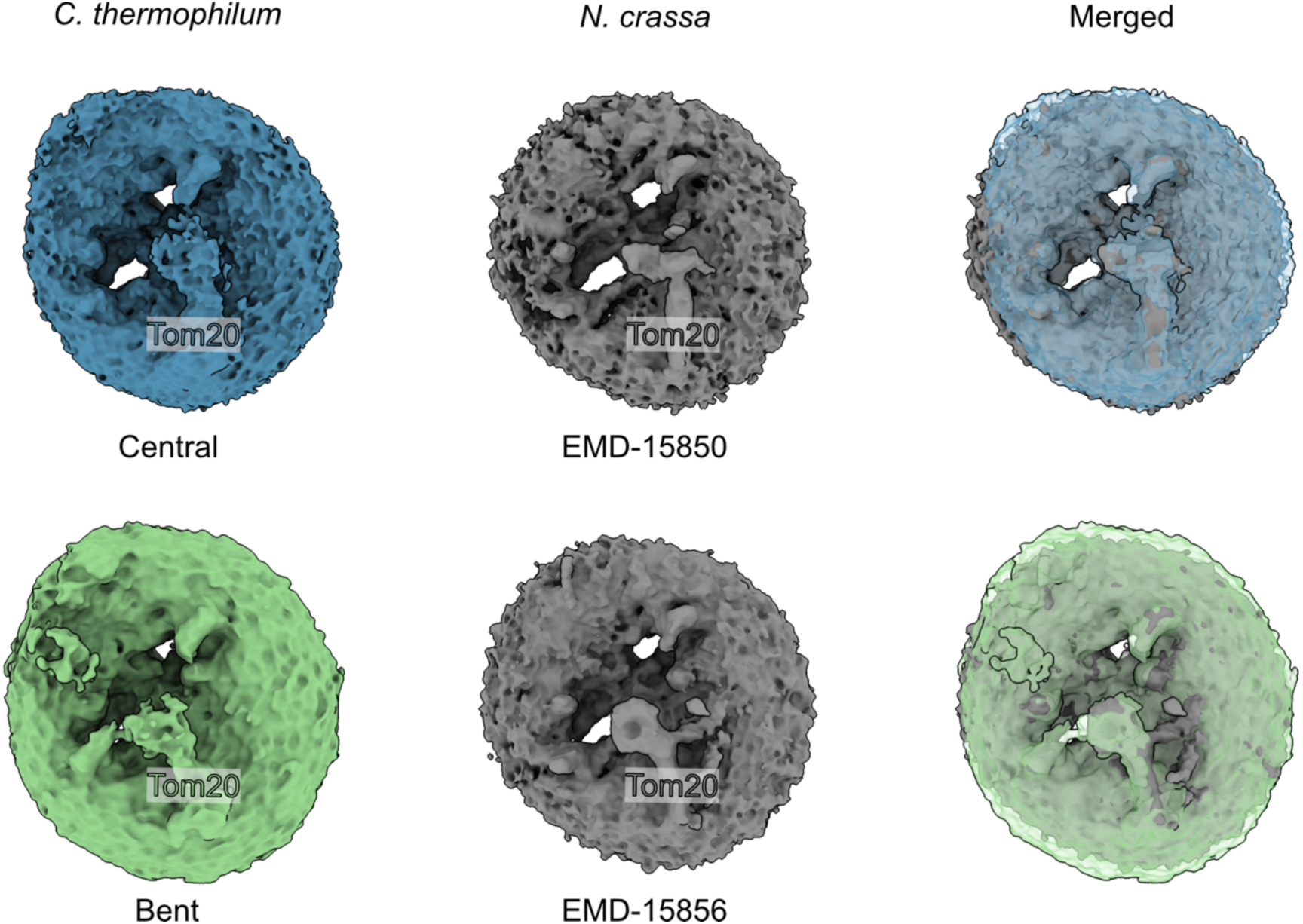
Conformational differences of Tom20 between *C. thermophilum* and *N. crassa*. Two conformations from *C. thermophilum* (Figure 4) are compared side-by-side and superposed to that of *N. crassa*. The contour level for each cryoEM map was set in UCSF Chimerax (1) so that Tom20 (labeled) would be visible.

**Figure S17.**
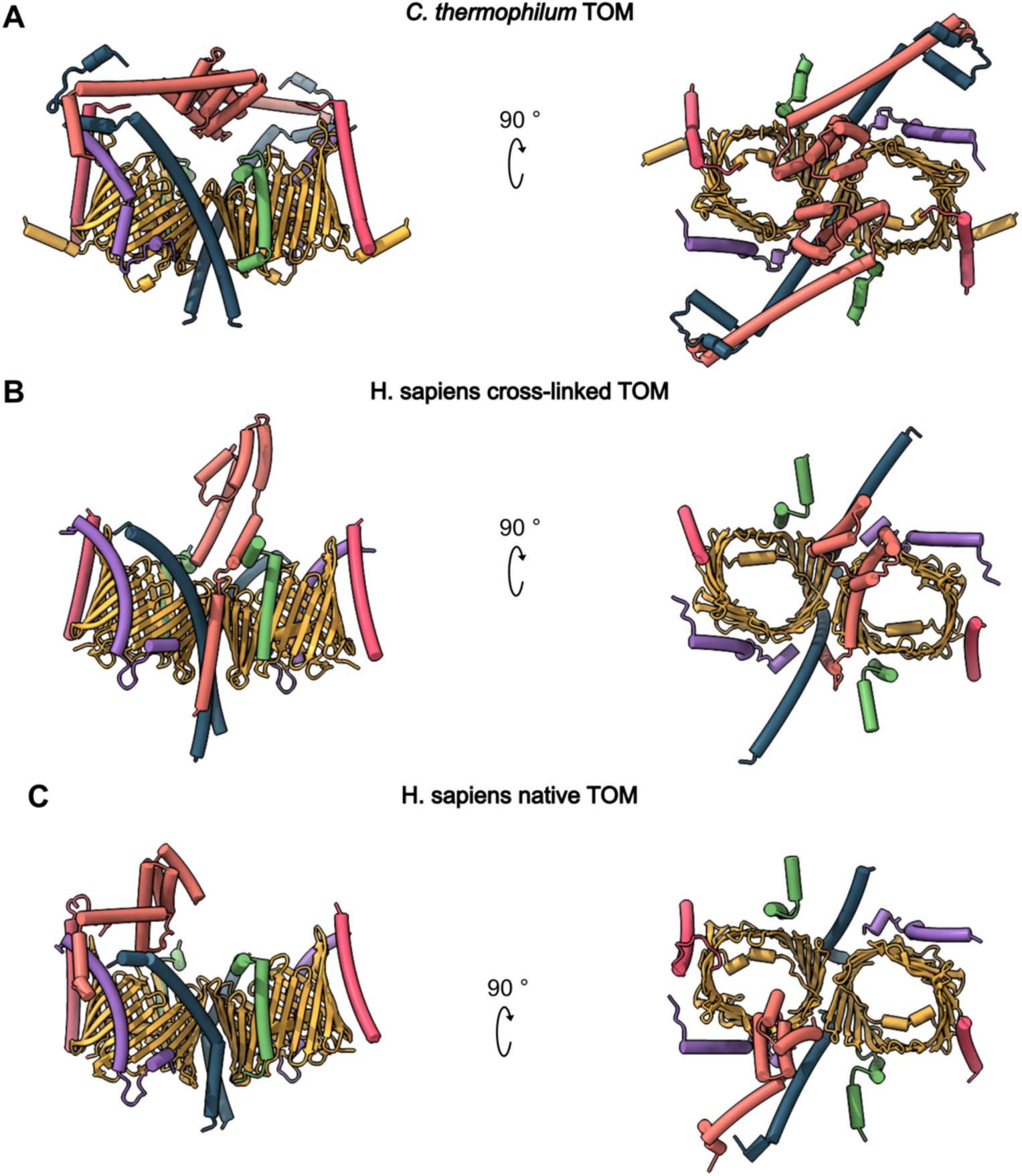
Positions of Tom20 in *H. sapiens* compared to *C. thermophilum.*. Our *C. thermophilum* TOM holo complex (**A**) compared to the cross-linked (**B**) human TOM (PDB: 8XVA) and (**C**) TOM subunits in the human TOM-PINK1-VDAC complex (PDB: 9EIH). The colours of TOM holo subunits are: Tom40, yellow; Tom22, blue; Tom20, orange; Tom5, Tom6 and Tom7, red, green and purple.

**Table S1.**
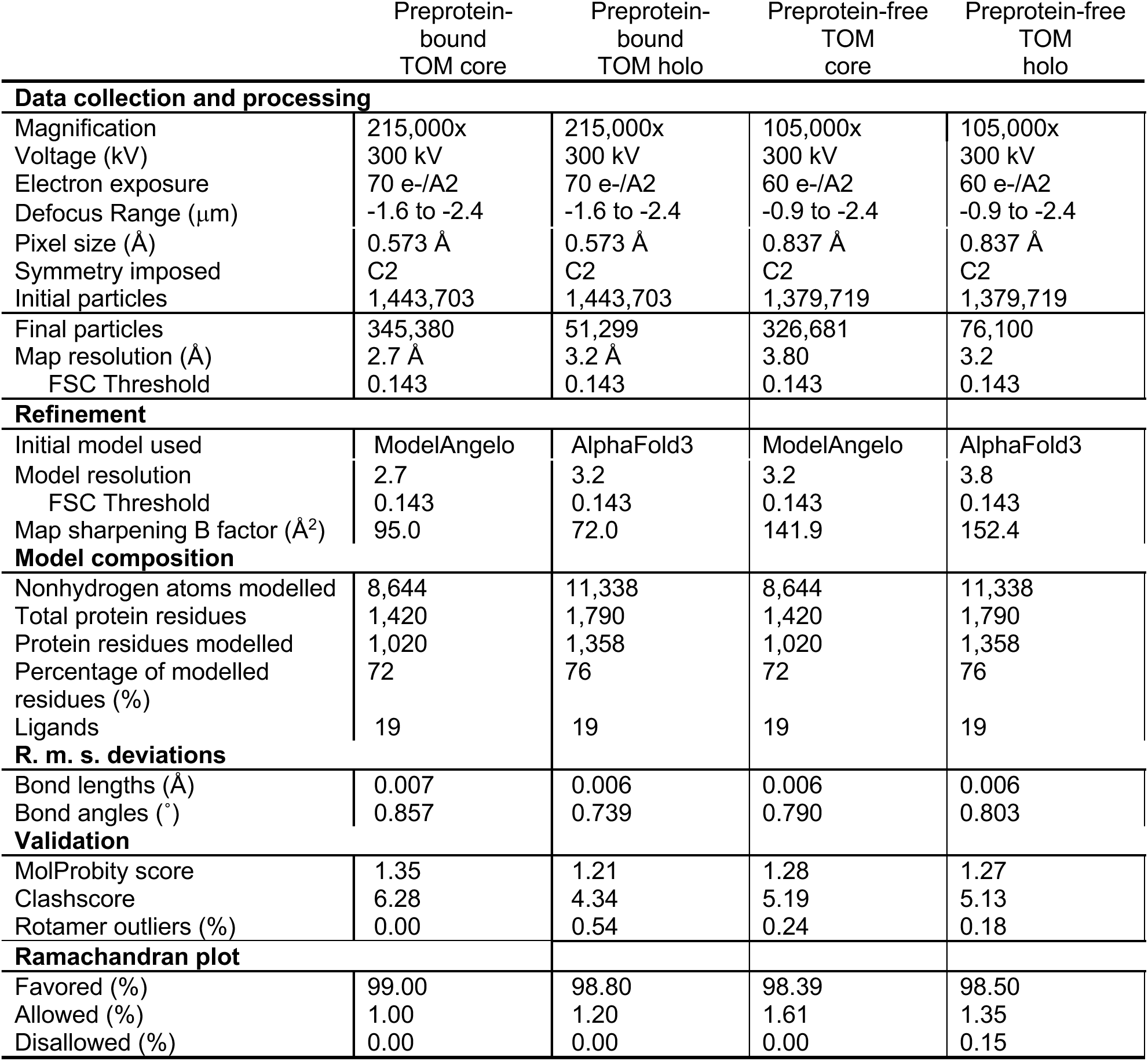
CryoEM data collection, refinement and validation of the *C. thermophilum* TOM core complex.

|  | Preprotein-bound<br>TOM core | Preprotein-bound<br>TOM holo | Preprotein-free<br>TOM<br>core | Preprotein-free<br>TOM<br>holo |
| --- | --- | --- | --- | --- |
| **Data collection and processing** |  |  |  |  |
| Magnification | 215,000x | 215,000x | 105,000x | 105,000x |
| Voltage (kV) | 300 kV | 300 kV | 300 kV | 300 kV |
| Electron exposure | 70 e-/Å<sup>2</sup> | 70 e-/Å<sup>2</sup> | 60 e-/Å<sup>2</sup> | 60 e-/Å<sup>2</sup> |
| Defocus Range (μm) | -1.6 to -2.4 | -1.6 to -2.4 | -0.9 to -2.4 | -0.9 to -2.4 |
| Pixel size (Å) | 0.573 Å | 0.573 Å | 0.837 Å | 0.837 Å |
| Symmetry imposed | C2 | C2 | C2 | C2 |
| Initial particles | 1,443,703 | 1,443,703 | 1,379,719 | 1,379,719 |
| Final particles | 345,380 | 51,299 | 326,681 | 76,100 |
| Map resolution (Å) | 2.7 Å | 3.2 Å | 3.80 | 3.2 |
| FSC Threshold | 0.143 | 0.143 | 0.143 | 0.143 |
| **Refinement** |  |  |  |  |
| Initial model used | ModelAngelo | AlphaFold3 | ModelAngelo | AlphaFold3 |
| Model resolution | 2.7 | 3.2 | 3.2 | 3.8 |
| FSC Threshold | 0.143 | 0.143 | 0.143 | 0.143 |
| Map sharpening B factor (Å<sup>2</sup>) | 95.0 | 72.0 | 141.9 | 152.4 |
| **Model composition** |  |  |  |  |
| Nonhydrogen atoms modelled | 8,644 | 11,338 | 8,644 | 11,338 |
| Total protein residues | 1,420 | 1,790 | 1,420 | 1,790 |
| Protein residues modelled | 1,020 | 1,358 | 1,020 | 1,358 |
| Percentage of modelled<br>residues (%) | 72 | 76 | 72 | 76 |
| Ligands | 19 | 19 | 19 | 19 |
| **R. m. s. deviations** |  |  |  |  |
| Bond lengths (Å) | 0.007 | 0.006 | 0.006 | 0.006 |
| Bond angles (°) | 0.857 | 0.739 | 0.790 | 0.803 |
| **Validation** |  |  |  |  |
| MolProbity score | 1.35 | 1.21 | 1.28 | 1.27 |
| Clashscore | 6.28 | 4.34 | 5.19 | 5.13 |
| Rotamer outliers (%) | 0.00 | 0.54 | 0.24 | 0.18 |
| **Ramachandran plot** |  |  |  |  |
| Favored (%) | 99.00 | 98.80 | 98.39 | 98.50 |
| Allowed (%) | 1.00 | 1.20 | 1.61 | 1.35 |
| Disallowed (%) | 0.00 | 0.00 | 0.00 | 0.15 |

## References

1. S. Rath et al., MitoCarta3.0: an updated mitochondrial proteome now with sub-organelle localization and pathway annotations. Nucleic Acids Res 49, D1541–D1547 (2021).

2. N. Wiedemann, N. Pfanner, Mitochondrial Machineries for Protein Import and Assembly. Annu Rev Biochem 86, 685–714 (2017).

3. U. Ahting et al., The TOM core complex: the general protein import pore of the outer membrane of mitochondria. J Cell Biol 147, 959–968 (1999).

4. P. J. Dekker et al., Preprotein translocase of the outer mitochondrial membrane: molecular dissection and assembly of the general import pore complex. Mol Cell Biol 18, 6515–6524 (1998).

5. J. C. Young, N. J. Hoogenraad, F. U. Hartl, Molecular chaperones Hsp90 and Hsp70 deliver preproteins to the mitochondrial import receptor Tom70. Cell 112, 41–50 (2003).

6. A. Chacinska, C. M. Koehler, D. Milenkovic, T. Lithgow, N. Pfanner, Importing mitochondrial proteins: machineries and mechanisms. Cell 138, 628–644 (2009).

7. K. Model, C. Meisinger, W. Kuhlbrandt, Cryo-electron microscopy structure of a yeast mitochondrial preprotein translocase. J Mol Biol 383, 1049–1057 (2008).

8. T. Bausewein et al., Cryo-EM Structure of the TOM Core Complex from Neurospora crassa. Cell 170, 693–700 e697 (2017).

9. Z. Guan et al., Structural insights into assembly of human mitochondrial translocase TOM complex. Cell Discov 7, 22 (2021).

10. K. Tucker, E. Park, Cryo-EM structure of the mitochondrial protein-import channel TOM complex at near-atomic resolution. Nat Struct Mol Biol 26, 1158–1166 (2019).

11. W. Wang et al., Atomic structure of human TOM core complex. Cell Discov 6, 67 (2020).

12. Y. Araiso et al., Structure of the mitochondrial import gate reveals distinct preprotein paths. Nature 575, 395–401 (2019).

13. A. Periasamy et al., Structure of an ex vivoDrosophila TOM complex determined by single-particle cryoEM. IUCrJ 12, 49–61 (2025).

14. P. Ornelas et al., Two conformations of the Tom20 preprotein receptor in the TOM holo complex. Proc Natl Acad Sci U S A 120, e2301447120 (2023).

15. J. Su et al., Structure of the intact Tom20 receptor in the human translocase of the outer membrane complex. PNAS Nexus 3, pgae269 (2024).

16. J. Su et al., Structural basis of Tom20 and Tom22 cytosolic domains as the human TOM complex receptors. Proc Natl Acad Sci U S A 119, e2200158119 (2022).

17. S. Callegari et al., Structure of human PINK1 at a mitochondrial TOM-VDAC array. Science 10.1126/science.adu6445, eadu6445 (2025).

18. X. Gao et al., Crystal structure of SARS-CoV-2 Orf9b in complex with human TOM70 suggests unusual virus-host interactions. Nat Commun 12, 2843 (2021).

19. S. Kreimendahl, J. Rassow, The Mitochondrial Outer Membrane Protein Tom70-Mediator in Protein Traffic, Membrane Contact Sites and Innate Immunity. Int J Mol Sci 21 (2020).

20. Y. Zhang et al., Structure of the mitochondrial TIM22 complex from yeast. Cell Res 31, 366–368 (2021).

21. L. Qi et al., Cryo-EM structure of the human mitochondrial translocase TIM22 complex. Cell Res 31, 369–372 (2021).

22. S. I. Sim, Y. Chen, D. L. Lynch, J. C. Gumbart, E. Park, Structural basis of mitochondrial protein import by the TIM23 complex. Nature 621, 620–626 (2023).

23. X. Zhou et al., Molecular pathway of mitochondrial preprotein import through the TOM-TIM23 supercomplex. Nat Struct Mol Biol 30, 1996–2008 (2023).

24. M. Kornprobst et al., Architecture of the 90S Pre-ribosome: A Structural View on the Birth of the Eukaryotic Ribosome. Cell 166, 380–393 (2016).

25. J. Abramson et al., Addendum: Accurate structure prediction of biomolecular interactions with AlphaFold 3. Nature 636, E4 (2024).

26. T. Saitoh et al., Tom20 recognizes mitochondrial presequences through dynamic equilibrium among multiple bound states. EMBO J 26, 4777–4787 (2007).

27. Y. Abe et al., Structural basis of presequence recognition by the mitochondrial protein import receptor Tom20. Cell 100, 551–560 (2000).

28. T. Shiota et al., Molecular architecture of the active mitochondrial protein gate. Science 349, 1544–1548 (2015).

29. S. Schmitt et al., Role of Tom5 in maintaining the structural stability of the TOM complex of mitochondria. J Biol Chem 280, 14499–14506 (2005).

30. A. I. de Kroon, D. Dolis, A. Mayer, R. Lill, B. de Kruijff, Phospholipid composition of highly purified mitochondrial outer membranes of rat liver and Neurospora crassa. Is cardiolipin present in the mitochondrial outer membrane? Biochim Biophys Acta 1325, 108–116 (1997).

31. M. H. Schuler et al., Phosphatidylcholine affects the role of the sorting and assembly machinery in the biogenesis of mitochondrial beta-barrel proteins. J Biol Chem 290, 26523–26532 (2015).

32. T. Becker et al., Role of phosphatidylethanolamine in the biogenesis of mitochondrial outer membrane proteins. J Biol Chem 288, 16451–16459 (2013).

33. N. Gebert et al., Mitochondrial cardiolipin involved in outer-membrane protein biogenesis: implications for Barth syndrome. Curr Biol 19, 2133–2139 (2009).

34. S. Nussberger, R. Ghosh, S. Wang, New insights into the structure and dynamics of the TOM complex in mitochondria. Biochem Soc Trans 52, 911–922 (2024).

35. D. Gessmann et al., Improving the resistance of a eukaryotic beta-barrel protein to thermal and chemical perturbations. J Mol Biol 413, 150–161 (2011).

36. D. Gessmann et al., Structural elements of the mitochondrial preprotein-conducting channel Tom40 dissolved by bioinformatics and mass spectrometry. Biochim Biophys Acta 1807, 1647–1657 (2011).

37. K. Yamano et al., Tom20 and Tom22 share the common signal recognition pathway in mitochondrial protein import. J Biol Chem 283, 3799–3807 (2008).

38. T. Shiota, H. Mabuchi, S. Tanaka-Yamano, K. Yamano, T. Endo, In vivo protein-interaction mapping of a mitochondrial translocator protein Tom22 at work. Proc Natl Acad Sci U S A 108, 15179–15183 (2011).

39. A. Schneider, Evolution and diversification of mitochondrial protein import systems. Curr Opin Cell Biol 75, 102077 (2022).

40. K. A. Rimmer et al., Recognition of mitochondrial targeting sequences by the import receptors Tom20 and Tom22. J Mol Biol 405, 804–818 (2011).

41. S. Rout et al., Determinism and contingencies shaped the evolution of mitochondrial protein import. Proc Natl Acad Sci U S A 118 (2021).

42. N. Kellner, E. Hurt, Transformation of Chaetomium thermophilum and Affinity Purification of Native Thermostable Protein Complexes. Methods Mol Biol 2502, 35–50 (2022).

43. E. Laube, J. Meier-Credo, J. D. Langer, W. Kuhlbrandt, Conformational changes in mitochondrial complex I of the thermophilic eukaryote Chaetomium thermophilum. Sci Adv 8, eadc9952 (2022).

44. K. P. Kunkele et al., The preprotein translocation channel of the outer membrane of mitochondria. Cell 93, 1009–1019 (1998).

45. J. Zivanov et al., New tools for automated high-resolution cryo-EM structure determination in RELION-3. Elife 7 (2018).

46. A. Punjani, J. L. Rubinstein, D. J. Fleet, M. A. Brubaker, cryoSPARC: algorithms for rapid unsupervised cryo-EM structure determination. Nat Methods 14, 290–296 (2017).

47. T. Wagner et al., SPHIRE-crYOLO is a fast and accurate fully automated particle picker for cryo-EM. Commun Biol 2, 218 (2019).

48. T. Bepler et al., Positive-unlabeled convolutional neural networks for particle picking in cryo-electron micrographs. Nat Methods 16, 1153–1160 (2019).

49. P. B. Rosenthal, R. Henderson, Optimal determination of particle orientation, absolute hand, and contrast loss in single-particle electron cryomicroscopy. J Mol Biol 333, 721–745 (2003).

50. A. Punjani, D. J. Fleet, 3D variability analysis: Resolving continuous flexibility and discrete heterogeneity from single particle cryo-EM. J Struct Biol 213, 107702 (2021).

51. E. F. Pettersen et al., UCSF ChimeraX: Structure visualization for researchers, educators, and developers. Protein Sci 30, 70–82 (2021).

52. K. Jamali et al., Automated model building and protein identification in cryo-EM maps. Nature 628, 450–457 (2024).

53. P. Emsley, B. Lohkamp, W. G. Scott, K. Cowtan, Features and development of Coot. Acta Crystallogr D Biol Crystallogr 66, 486–501 (2010).

54. T. I. Croll, ISOLDE: a physically realistic environment for model building into low-resolution electron-density maps. Acta Crystallogr D Struct Biol 74, 519–530 (2018).

55. D. Liebschner et al., Macromolecular structure determination using X-rays, neutrons and electrons: recent developments in Phenix. Acta Crystallogr D Struct Biol 75, 861–877 (2019).

## SI References

1. E. F. Pettersen et al., UCSF ChimeraX: Structure visualization for researchers, educators, and developers. Protein Sci 30, 70–82 (2021).

2. J. Abramson et al., Addendum: Accurate structure prediction of biomolecular interactions with AlphaFold 3. Nature 636, E4 (2024).

